# Antarctic Soil Auxiliarome: unraveling the pan-auxiliary metabolic genes catalogue in a transect across different ice-free regions of Antarctica

**DOI:** 10.1101/2025.10.13.682075

**Authors:** Sergio Sanchez-Carrillo, Rafael Gonzalez-Serrano, María del Carmen Fernández-Moyano, S. Craig Cary, Ian R. McDonald, Antonio Alcamí

**Affiliations:** Centro de Biología Molecular Severo Ochoa, UAM-CSIC, Madrid, Spain; Department of Physiology, Genetics and Microbiology. Universidad de Alicante, Alicante, Spain; International Center for Terrestrial Antarctic Research, University of Waikato, Hamilton, New Zealand; Te Aka Matuatua-School of Science, Te Whare Wānanga o Waikato-University of Waikato, Hamilton, New Zealand

**Keywords:** auxiliarome, metavirome, auxiliary metabolic gene, transantarctic mountains, Antarctica, ice-free region

## Abstract

The Transantarctic Mountains host a significant portion of Antarctica’s ice-free soils and support diverse microbial communities, but the role of viruses in these extreme ecosystems remains poorly understood. To address this gap, we conducted the first comprehensive analysis of both RNA and DNA soil viruses across ten locations along a regional-scale latitudinal transect. This study revealed high viral diversity, including the first description of 18 previously unreported viral families in Antarctic soils, alongside significant local viral endemicity. Elevation emerged as the primary driver of viral diversity, while distance to the coast explained the distribution of auxiliary metabolic genes (AMGs), and distance to the sea influenced the metabolic pathways associated with these AMGs. The concept of the "auxiliarome," a pan-AMG catalogue at the community level, underscores the critical role of AMGs in engineering host metabolism. These genes contribute to the metabolism of cofactors and vitamins, amino acids, carbohydrates, and sulfur, as well as ecologically significant traits such as bacterial restriction-modification systems or antibiotic production and resistance. This study expands our understanding of Antarctic soil viruses and highlights their ecological importance in shaping microbial communities and biogeochemical processes in extreme environments.

## Introduction

Ice-free regions of Antarctica cover less than 0.5% of the continent’s surface, predominantly located in the Transantarctic Mountains (TAM), with the McMurdo Dry Valleys being one of the most representative areas [1]. The TAM, a mountain range stretching 3,200 km from Cape Adare in northern Victoria Land to Coats Land, divides the continent into East and West Antarctica, reaching its highest elevation at Mt. Kirkpatrick (4,528 m) [2]. Opposite the coast of Victoria Land on Ross Island, Mt. Erebus rises 3,794 m above the Ross Sea and is the southernmost active volcano on Earth [3], hosting exposed ice-free geothermal soils [4]. TAM soils are classified as permafrost-affected soils, belonging to the Gelisol order [5], while Mt. Erebus soils are lithosols [4]. These ice-free regions present numerous environmental gradients, including elevation, climate, latitude, and geological diversity, which create a wide range of habitats and support highly diverse microbial communities [6–11].

Despite the extreme conditions of Antarctic ice-free soils—characterized by high UV radiation, low water and nutrient availability, high salinity, and freezing temperatures—these soils harbor a rich and diverse microbial ecosystem [6–11]. Representatives of the three domains of life (Bacteria, Archaea, and Eukarya) coexist with viruses, which infect and interact with them. The interplay between biotic and abiotic factors shapes the structure and function of these unique ecosystems [6, 9, 12], although these dynamics remain poorly understood.

Viruses are the most abundant and genetically diverse biological entities on Earth [13], exerting a profound ecological influence on microbial ecosystems. They regulate microbial populations through parasitism and contribute to biogeochemical nutrient cycling via the viral shunt [14–22]. This process prevents particulate organic matter from ascending to higher trophic levels, recycling it into dissolved organic matter that re-enters the microbial loop [19]. Viruses drive microbial evolution through their roles as evolutionary catalysts, shaping host anti-viral responses in an ongoing arms race [23–29] and facilitating horizontal gene transfer across species and families [30–33]. Additionally, viruses engineer host metabolism by exchanging AMGs with their hosts [34–45]. These AMGs, which are host-derived but virus-encoded, correspond to non-replicative functions and are incorporated into viral genomes during infection. If these genes confer fitness advantages, such as enhanced metabolic capabilities, then they are retained [46]. The use of AMGs by different viral communities has been associated with an impact on critical biogeochemical cycles, influencing the cycling of carbon [42, 44], nitrogen [37, 39], sulfur [36], and phosphorus [34].

Although soil viruses play a critical role in controlling microbial dynamics, influencing host metabolic states, and driving biogeochemical processes, they remain underexplored in Antarctic ecosystems. Challenges such as low biomass and restricted accessibility have limited research efforts, leaving significant gaps in our understanding of viral communities in these extreme environments.

To address this, we analyzed ten metaviromes collected across a meso-scale latitudinal transect spanning the TAM and Mt. Erebus. This study represents the first comprehensive examination of both DNA and RNA viruses in Antarctic soils at this geographical scale. Additionally, we introduce the concept of the soil auxiliarome, a pan-AMG catalogue that integrates the biochemical potential of viruses to modulate host metabolism and contribute to ecosystem-level processes. By exploring the diversity of viral communities and their functional contributions, this work provides valuable insights into the ecological roles of viruses in Antarctic ice-free soils.

## Material and methods

### Site description and sample collection

Soil samples were collected in several ice-free regions from different locations in the TAM. We also included a sample from Mt. Erebus (ERE) (Figure 1b). The Northern Victoria Land samples were collected by Craig Cary. The Mt Erebus/Ross Island samples and the McMurdo Dry Valley samples were collected by Craig Cary and Ian McDonald. The Central TAM samples were collected by Prof Hugh Morgan and Prof Ian Hogg (Waikato University).

**Figure 1.**
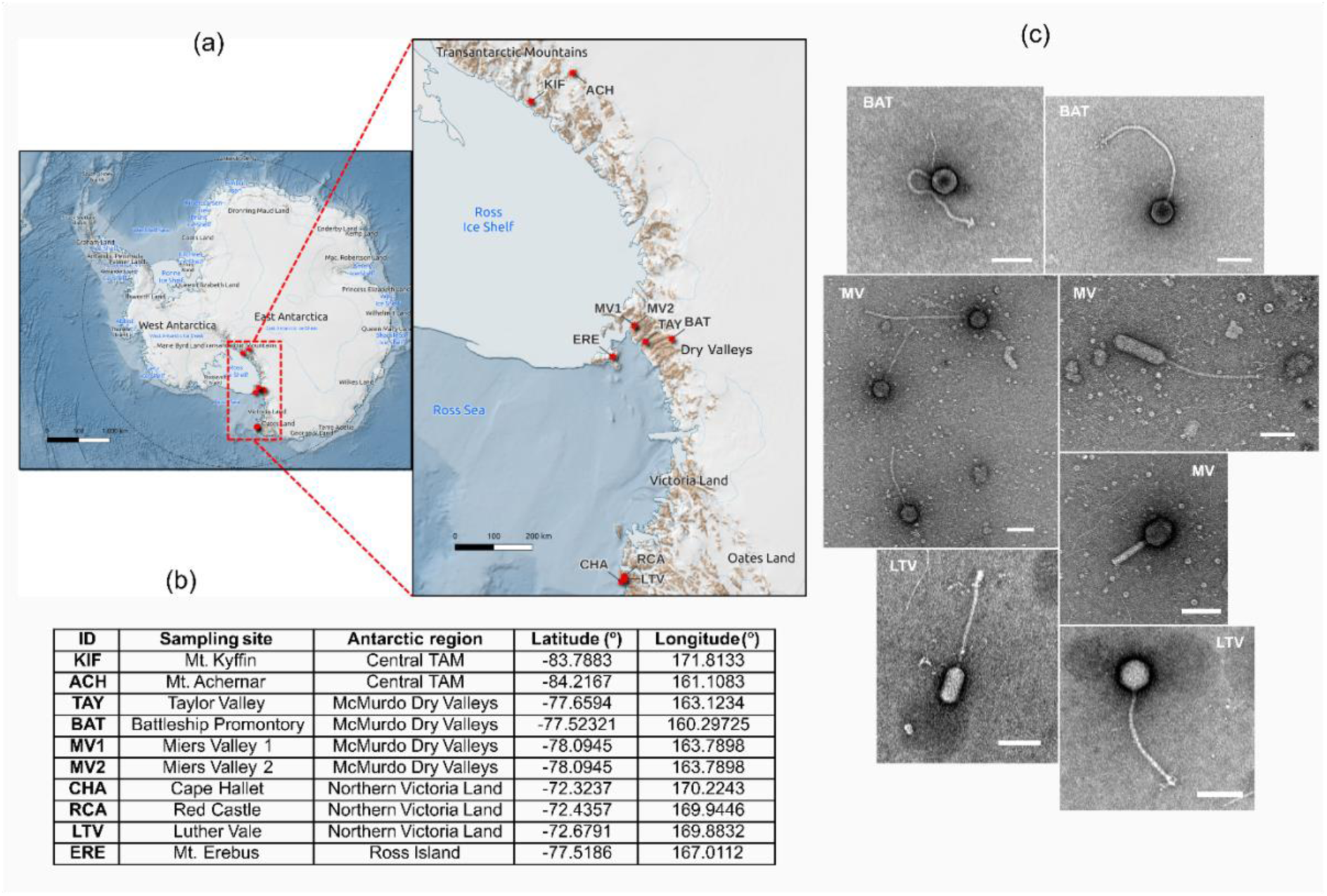
**(a)** General (left) and zoomed-in (right) maps of Antarctic ice-free soil sampling sites. Red asterisks indicate sampling locations, and brown shading denotes ice-free areas. See sample identifiers in table of panel (b). **(b)** Antarctic soil sampling locations, including location names, identifiers, Antarctic regions, and geographical latitude and longitude. **(c)** Transmission electron microscopy images of *Caudaviricetes* viruses identified at Battleship Promontory (BAT), Miers Valley (MV) and Luther Vale (LTV). Scale bar: 100 nm.

The location of the sampling sites is shown in Figure 1a, and the main characteristics of the sites in Table S1. All samples form a latitudinal transect that spans approximately 1,200 km, from Cape Hallet (CHA) at 72.3236°S to Mt. Achernar (ACH) at 84.217°S.

Sampling of the TAM [11] and ERE soils [47] has been described previously. In all cases samples were collected by first discarding the top centimetre of soil, in particular the pavement pebbles, and collecting the next 2 - 3 cm of soil using a sterile spatula. Samples were placed into sterile 50 ml tubes or Whirl-Paks (Nasco International, Fort Atkinson, WI, USA) and frozen for transport back to the University of Waikato (Hamilton, New Zealand) for analysis.

### Sample processing and viral purification

Soil samples were thawed in a laminar flow cabinet, and 30 g of each was transferred into a 50 ml falcon tube using a sterile spatula. To homogenize the samples, a modified SM buffer (50 mM Tris-HCl, 8 mM MgCl_2_, 100 mM NaCl, pH 7.5) was added to fill each tube and then left at 4 °C overnight. To remove the eukaryotic and prokaryotic cells from the sample and retain the remaining viral fraction, the tubes were agitated by vortexing for 20 minutes at 4° C and centrifuged two times at 3,000 x *g* for 20 minutes at 4° C. The remaining supernatant was filtered using 0.45 µm syringe filters with Supor membrane (PALL) in order to preserve large viral particles as well. Since the use of this filter pore size might imply the presence of some remaining prokaryotic cells in the samples, a 16S rRNA PCR assay was performed on all samples to detect possible prokaryotic DNA contamination (see below). Finally, samples were concentrated to a volume of ∼250 µl using 100 kDa Amicon centrifugal filter units. A 20 µl aliquot of the concentrate was saved for transmission electron microscopy (TEM) analysis. The laminar flow cabinet and part of the laboratory material used during the process was cleaned with DNA-ExitusPlus (PanReac AppliChem) and 0.1 N HCl to avoid DNA cross-contamination. Throughout the entire process, the appropriate personal protective equipment for working with this type of sample (double gloves sealed with paper tape, disposable sleeves, masks, etc) was used.

### Transmission electron microscopy

Samples were adsorbed for 3 min onto ionized Collodion/carbon coated copper grids and negatively stained with 2% aqueous uranyl acetate for 30 seconds. Grids were examined in a JEM1400 Flash transmission electron microscope (Jeol, Japan) at an accelerating voltage of 100kV and images were taken with a OneView CMOS 4Kx4K camera (Gatan, Pleasanton, CA USA) at the Electron Microscopy Service of the Centro de Biología Molecular Severo Ochoa (CBM, CSIC), Madrid.

### Viral nucleic acid extraction, amplification and sequencing

Prior to extraction of viral nucleic acids, a 16S rRNA PCR assay was performed on all samples using the universal primers 27F and 1492R for detection of possible prokaryotic DNA contamination. Then, samples were treated with the endonuclease Benzonase to remove all free forms of microbial DNA and RNA (single stranded, double stranded, linear and circular) present outside viral capsids. Briefly, 0.5 µl of Benzonase (125U) and 20 µl of a buffer containing 50 mM Tris-HCl and 2 mM MgCl_2_ (pH 8) were added to 180 µl of the sample and incubated for 30 min at 37° C. Then, 4 µl of 0.5 M EDTA (pH 8) was added at room temperature to stop the reaction. Viral nucleic acids (DNA and RNA) were extracted using the QIAamp MinElute Virus Spin Kit (QIAGEN) and the nucleic acids obtained were eluted in 50 µl of nuclease free water. DNA and RNA concentrations were determined separately using a Quantus Fluorometer (Promega) before storage at −80° C.

Due to the low viral biomass obtained in this type of environmental sample, the resulting viral DNA was amplified for 1.5 h with the enzyme Phi29 DNA polymerase and random hexamer primers by multiple displacement amplification (MDA) [48] using the GenomiPhi V3 Ready-To-Go DNA Amplification Kit (Cytiva). The resulting viral RNA was amplified by sequence - independent single-primer amplification (SISPA) [49, 50]. Briefly, the SuperScript IV First-Strand Synthesis System kit (Invitrogen) was used to synthesize first-strand cDNA from RNA using random primers FR20-12N and FR20-20T (FR20: 5’-GCCGGAGCTCTGCAGATATC-3’). Then the Klenow fragment enzyme (New England Biolabs) was used to convert cDNA into dsDNA using primer FR20-12N. Finally, PCR amplification with the dsDNA obtained in the previous step was performed with the enzyme Q5 High-Fidelity DNA Polymerase (New England Biolabs) (35 cycles) and random primers FR20, N-FR20, 2N-FR20 and 3N-FR20.

The DNA concentration of all amplified samples was measured with the Quantus Fluorometer (Promega) before storage at −20° C and DNA integrity of each sample was analyzed by agarose gel electrophoresis.

Before sequencing, all DNA of samples amplified by MDA were purified using the QIAquick PCR Purification kit (QIAGEN). And all DNA obtained with samples amplified by SISPA were purified using the CleanNGS kit (CleanNA) adding 1.8x the sample volume of CleanNGS reagent to each sample to remove DNA fragments smaller than 100 bp and retain the larger fragments. This is an efficient DNA purification kit based on paramagnetic beads technology. Finally, all samples were sequenced with Illumina NextSeq2000 (2×150 bp paired-end reads) at the Centre for Genomic Regulation (CRG), Barcelona, Spain.

### Computational analysis

Sequenced reads were quality checked using FastQC 0.12.0 (https://www.bioinformatics.babraham.ac.uk/projects/fastqc/) and multiQC 1.21 [51] and quality trimmed and filtered using Trimmomatic 0.39 [52] with parameters ILLUMINACLIP:TruSeq3-PE:2:30:10 LEADING:3 TRAILING:3 SLIDINGWINDOW:3:15 MINLEN:36 Assembly of the reads was performed using MEGAHIT 1.2.9 [53] with default parameters and --presets meta-large, for dealing with complex metagenomes. The assembled contigs were analyzed with VirSorter2 2.2.4 [54], VIBRANT 1.2.1 [55] and DeepVirFinder 1.0 [56] for viral sequence identification, combining the different sets obtained with each method for further analysis. Viral binning was performed with VRhyme 1.1.0 [57] keeping those contigs longer or equal to 2,000 bp and using the parameter --method composite to enable the scaffold dereplication combining overlapping scaffolds into composite contigs. To perform the quantification of the binned contigs, reads were mapped back to the contigs using CoverM 0.7.0 (https://github.com/wwood/CoverM) with the following parameters: contig (contig mode), --min-covered-fraction 0.95, --min-read-percent-identity 0.95 and --min-read-aligned-percent 0.95. The last 3 parameters were used to minimise spurious mappings that could falsely inflate abundances. Contigs quantified with a value of zero were removed. Quantification was done using TPM (transcripts per million).

Contigs were taxonomically annotated using geNomad 1.8.0 [58], with the ICTV taxonomy VMR 19 (https://ictv.global/vmr) in end-to-end mode with default parameters, keeping the contigs with virus_score ≥ 0.7 and deleting those annotated as plasmid with at least one plasmid hallmark gene.

Host prediction was performed using iPHoP [59] and vpf-tools [60] at the genus level (membership_ratio ≥ 0.5 and confidence_score ≥ 0.5), using the taxonomy for prokaryotes contained in the GTDB database 202 [61]. This annotation was manually curated for those contigs annotated at the family level and modified with the host domain obtained in the Virus-Host Database (https://www.genome.jp/virushostdb) in case of inconsistencies. To annotate the bins in terms of taxonomy and predicted host taxonomy, a lowest common ancestor (LCA) approach was carried out. Briefly, for each bin, all contigs belonging to it were considered, using the taxonomy of most contigs (≥60%) for each taxonomic rank, and leaving it as "unknown" otherwise. Bins with human or mouse predicted hosts were considered contamination and therefore removed.

AMGs were detected with VIBRANT 1.2.1 [55] using as input the predicted viral contigs, and KEGG (Kyoto Encyclopedia of Genes and Genomes) ontology [62–64] was used at KO (KEGG Orthology) and Pathway levels. Online KEGG Mapper 5 [65,66] was used to produce a metabolic map of all the AMGs previously predicted.

The quality (completeness and contamination) of the binned contigs was analyzed with CheckV [67] and those considered ‘High-quality’ in the MIUViG [68] column were assigned as viral metagenome-assembled genomes (vMAGs), some of them considered ‘complete’ by CheckV. Same tool was employed to determine whether the binned contigs follow a lytic or a lysogenic cycle.

Network building and visualization was performed with Cytoscape 3.10.2 [69].

### Geographical analysis

All the bioclimatic variables used in this work were extracted from CHELSA database 2.1 [70] using the R package terra 1.7-71 [71] following [6]. The extracted bioclimatic variables were BIO1 (mean annual air temperature, °C), BIO2 (mean diurnal air temperature range, °C), BIO4 (temperature seasonality, °C/100), BIO5 (mean daily maximum air temperature of the warmest month, °C), BIO10 (mean daily mean air temperatures of the warmest quarter, °C), BIO12 (annual precipitation, kg m^-2^), BIO14 (precipitation in the driest month, kg m^-2^), BIO15 (precipitation seasonality, %), BIO17 (mean monthly precipitation in the driest quarter, kg m^-2^), BIO18 (mean monthly precipitation in the warmest quarter, kg m^-2^), and SWE (snow water equivalent, kg m^-2^). The elevation values extracted from the REMA digital elevation model (100 m DEMs) [72], the distance to the coast and to the sea for each sample point, and all the maps contained in this work were obtained using QGIS Desktop 3.22 (http://www.qgis.org) in Quantarctica [73].

### Statistical analysis

All calculations and plots were performed using the R programming language 4.3.3 [74] with packages vegan 2.6.4 [75] for biodiversity analysis, ComplexHeatmap 2.16.0 [76,77] for heatmaps, Rtsne 0.17 [78] for t-distributed Stochastic Neighbor Embedding (t-SNE), cluster 2.1.6 [79] for partition around medioids (PAM) clustering, UpSetR 1.4.0 [80] for intersecting sets, ggplot2 3.5.0 [81] for general plotting, ggforce 0.4.2 [82] for sankey plot and tidyverse 2.2.0 [83] for data manipulation.

To study the α-diversity, the community matrix in terms of bins was standardized employing the Hellinger method with decostand (vegan) and then used to calculate the Shannon and Simpson indexes with diversity (vegan). To calculate chao1 index the previous matrix was scaled to pseudo absolute frequencies using the minimum sample value and rounded to the next integer before using estimateR (vegan).

Rarefaction curves were obtained with rarecurve (vegan) using the previously calculated pseudoabsolute frequencies matrix and taking as subsampling size the minimum sum of rows of the matrix.

Environmental variables were standardized using decostand (vegan) before applying a redundancy analysis (RDA) with rda (vegan). Community matrix was standardized employing the Hellinger method with decostand (vegan). R^2^ and adjusted R^2^ were calculated with RsquareAdj (vegan). The significance of the RDA model, the environmental variables and the RDA axis were obtained using an overall permutation test (n=1000) with anova.cca (vegan). Only those environmental variables with p≤0.05 in the previous test were plotted in the RDA biplot and their coordinates were calculated using envfit (vegan) and scores (vegan). Forward selection was done from the simplest model (without variables) to the complete model (all variables) employing ordiR2step (vegan) and using adjusted R2 as the stopping criterion. Environmental variables selected in forward selection were considered as main drivers.

The t-SNE analysis was done with Rtsne (Rtsne) using the Jaccard distance calculated over the previously described standardized community matrix using vegdist (vegan). The PAM clustering was performed with the pam (cluster) function calculating the best silhouette width using silhouette (cluster). This analysis was performed also over the AMGs and the metabolic pathways they belong. Indicator species analysis (ISA) was performed using indicspecies (multipatt), using the previously derived PAM cluster assignments. Statistical significance was assessed by 1,000 permutations, and only individual clusters (no cluster combinations) were tested.

## Results

We analyzed nine soil metaviromes collected from three ice-free regions of the TAM— Northern Victoria Land, McMurdo Dry Valleys, and Central TAM—along with one metavirome from ERE, a volcano on Ross Island near the Dry Valleys (Figure 1a and 1b). The samples span Antarctic campaigns conducted between 2006 and 2020 (Table S1). These metaviromes provided a comprehensive dataset to explore viral diversity and their functional potential across diverse Antarctic soil environments. The dataset comprised both DNA and RNA viruses, providing a more comprehensive view of the virosphere in these extreme environments.

TEM images revealed different morphotypes of *Caudaviricetes* viruses in Battleship Promontory (BAT), Miers Valley (MV), and Luther Vale (LTV) samples (Figure 1c). These included siphoviral morphologies with their typical icosahedral capsids and long and flexible tails (BAT, MV left and LTV right); siphovirus-like phages with elongated capsids (MV upper right and LTV left); and phages with myoviral morphology (MV bottom right). These structural observations were consistent with the dominant viral groups identified by sequencing.

Sequencing of the DNA and RNA viromes yielded a total of 814,781,976 reads (average length: 150 bp), encompassing 122,217 Mbp of DNA (Table S2). After quality filtering, trimming (see Table S3), and assembly (Table S4), we obtained 66,430 metagenomic viral contigs (mVCs) clustered into 51,559 metagenomic viral bins (mVBs). Of these, 99.8% were DNA mVCs (66,274 contigs; 51,444 bins), and 0.2% were RNA mVCs (156 contigs; 115 bins). No RNA mVCs were detected in the ACH or ERE samples (Table S4). Most of the mVBs (72.5%) were singletons (comprising only one mVC each), with the remaining 27.5% comprising 3,383 multi-contig bins (mean = 5.4 mVCs per bin; SD = 10.0 mVCs). Table 1 provides a more comprehensive summary of the definitive viral assemblage statistics.

**Table 1:** Summary of viral statistics for Antarctic ice-free soils samples. Columns: (1) sample identifier (ACH: Mt. Achernar; BAT: Battleship Promontory; CHA: Cape Hallet; ERE: Mt. Erebus; KIF: Mt. Kyffin; LTV: Luther Vale; MV: Miers Valley; RCA: Red Castle; TAY: Taylor Valley); (2) viral type (DNA or RNA); (3) total number of contigs detected; (4) proportion of high-quality (HQ) contigs; (5) proportion of complete (C) contigs; (6) number of identified proviruses (P); (7) mean contig length (*L̅*); (8) N50 value; and (9) total number of bins. Percentages in columns 4 and 5 are calculated relative to the total contig count in column 3. Note that contig counts here may differ from those reported in the assembly output as these numbers refer to final contigs.

| Id | Type | Contigs | HQ (%) | C (%) | P (%) | $\bar{L}$ (bp) | N50 (bp) | Bins |
| --- | --- | --- | --- | --- | --- | --- | --- | --- |
| ACH | DNA | 491 | 1 (0.2) | 1 (0.2) | 9 (1.8) | 5,951.2 | 7,142 | 214 |
| BAT | DNA | 10,897 | 59 (0.5) | 33 (0.3) | 149 (1.4) | 5,134.1 | 5,968 | 7,990 |
| BAT | RNA | 19 | 3 (15.8) | 0 | 0 | 2,840.6 | 2,937 | 17 |
| CHA | DNA | 5,671 | 22 (0.4) | 9 (0.2) | 78 (1.4) | 5,158.3 | 5,960 | 3,568 |
| CHA | RNA | 7 | 0 | 0 | 0 | 3,024.3 | 3,005 | 7 |
| ERE | DNA | 286 | 0 | 0 | 2 (0.7) | 12,020.3 | 39,363 | 164 |
| KIF | DNA | 7,593 | 36 (0.5) | 18 (0.2) | 102 (1.3) | 4,493.7 | 4,846 | 5,604 |
| KIF | RNA | 58 | 8 (13.8) | 0 | 0 | 2,885.9 | 2,950 | 33 |
| LTV | DNA | 7,010 | 0 | 0 | 7 (0.1) | 4,578.1 | 4,823 | 5,668 |
| LTV | RNA | 23 | 0 | 0 | 0 | 2,871.2 | 2,823 | 22 |
| MV1 | DNA | 15,902 | 33 (0.2) | 21 (0.1) | 170 (1.1) | 4,156.8 | 4,346 | 13,101 |
| MV1 | RNA | 46 | 9 (19.6) | 0 | 0 | 2,988.6 | 3,017 | 35 |
| MV2 | DNA | 15,916 | 31 (0.2) | 20 (0.1) | 182 (1.1) | 4,128.6 | 4,280 | 13,342 |
| MV2 | RNA | 36 | 10 (27.8) | 0 | 0 | 3,098.6 | 3,065 | 29 |
| RCA | DNA | 9,005 | 26 (0.3) | 16 (0.2) | 133 (1.5) | 4,456.8 | 4,795 | 7,626 |
| RCA | RNA | 5 | 0 | 0 | 0 | 2,766.2 | 3,173 | 5 |
| TAY | DNA | 4,843 | 5 (0.1) | 0 | 42 (0.9) | 3,481.1 | 3,343 | 3,718 |
| TAY | RNA | 15 | 0 | 0 | 0 | 3,133.1 | 3,285 | 15 |

We identified 241 vMAGs, including 211 DNA sequences (range: 2.0–212.2 kbp; mean ± SD: 31.6 ± 27.2 kbp) and 30 RNA sequences (3.6–5.2 kbp; 4.0 ± 0.4 kbp). Most DNA vMAGs belonged to the class *Caudoviricetes* with no predicted host assigned, while RNA vMAGs were classified as *Leviviricetes*, also with unknown host. Of all DNA vMAGs, 116 sequences were classified as complete (2.0–132.5 kbp; 27.0 ± 22.9 kbp). The complete taxonomic annotation for all mVCs and mVBs is provided in Table S5.

### Taxonomic and Diversity Analysis

Viral bins distribution showed high endemicity, except between Miers Valley 1 (MV1) and Miers Valley 2 (MV2), both collected at the same sampling site (Figures 1a and 1b). Shannon α-diversity indices showed MV1 and MV2 as the most diverse, followed by BAT, LTV, and Red Castle (RCA) (Table 2). However, ACH and ERE showed the lowest diversity, probably indicating the presence of only a few taxa.

**Table 2:** Viral α-diversity indexes found in Antarctic soil samples, containing Shannon, Simpson, Chao1 and richness indexes obtained from mVBs.

| Sample Id | Shannon | Simpson | Chao1 | Richness |
| --- | --- | --- | --- | --- |
| ACH | 4.8764 | 0.9886 | 216.8 | 214 |
| BAT | 8.5278 | 0.9994 | 8,161.3 | 8,007 |
| CHA | 7.5318 | 0.9983 | 3,727.8 | 3,575 |
| ERE | 4.3947 | 0.9769 | 174.6 | 164 |
| KIF | 8.0440 | 0.9990 | 6,106.4 | 5,637 |
| LTV | 8.1871 | 0.9993 | 5,810.1 | 5,690 |
| MV1 | 9.0941 | 0.9997 | 13,171.2 | 13,136 |
| MV2 | 9.0973 | 0.9997 | 13,438.7 | 13,371 |
| RCA | 8.5745 | 0.9993 | 7,829.7 | 7,631 |
| TAY | 7.7892 | 0.9991 | 3,748.1 | 3,733 |

Rarefaction curves confirmed adequate sampling depth, as indicated by the curves tapering off and moving away from the initial exponential growth phase (Figure 2c). This suggests that most bins were successfully detected, minimizing underrepresentation of diversity.

**Figure 2:**
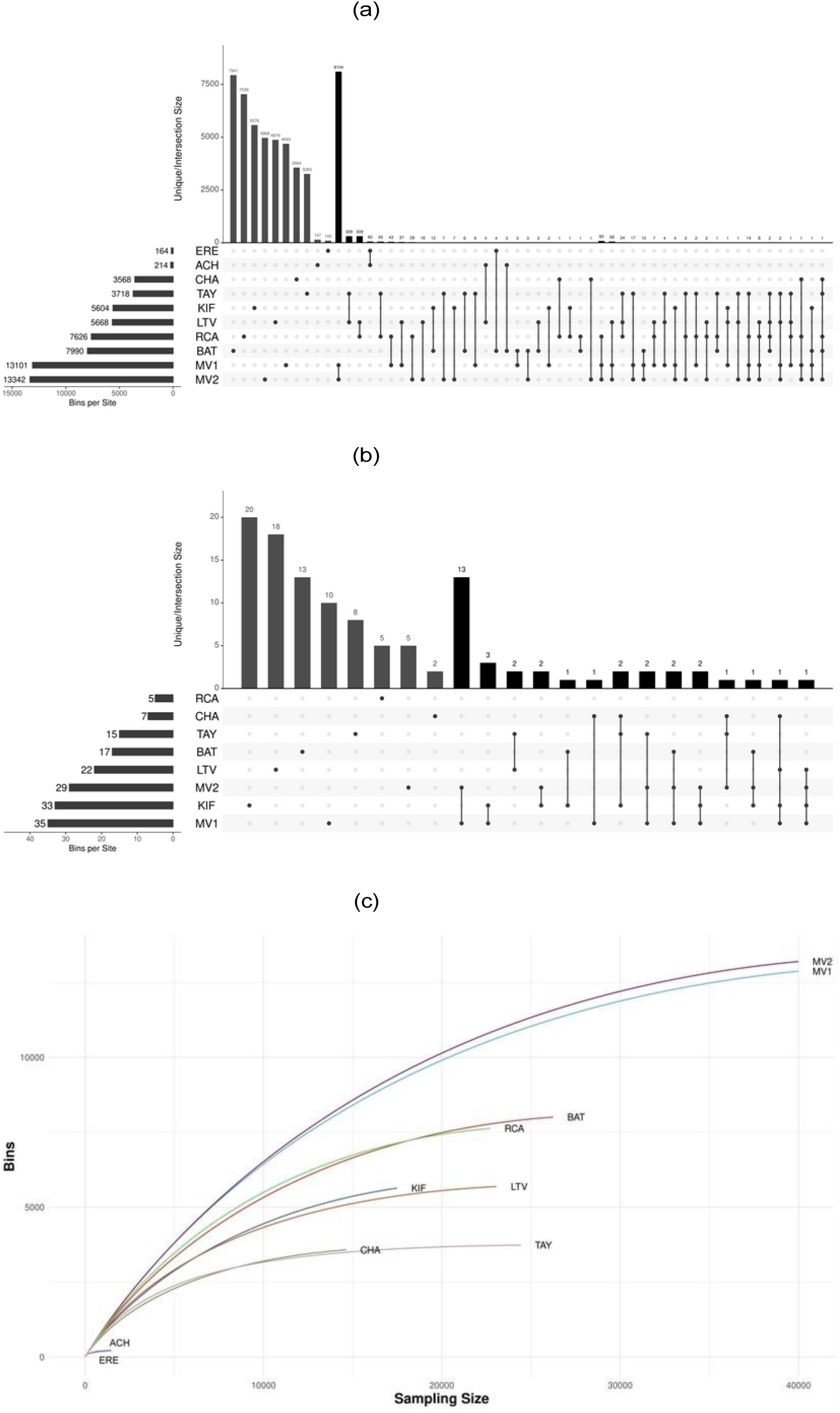
**(a-b)** UpSet plots showing the total number of mVBs per sample (black horizontal bars), unique mVBs per sample (gray vertical bars), and mVBs shared between samples (black vertical bars) for (a) DNA and (b) RNA bins. **(c)** Rarefaction curves for total mVBs (DNA and RNA), where "Bins" refers to the number of mVBs and "Sampling Size" corresponds to pseudo-absolute frequencies (i.e., pseudo reads) per sample.

The taxonomic distribution of mVBs in each sample revealed *Caudoviricetes* as the dominant viral DNA class in most samples, while RNA viruses, including *Leviviricetes* and *Duplopiviricetes*, constituted a smaller proportion (Figure 3); diversity patterns observed in this figure were consistent with those depicted in Figures 2a and 2b, where MV1 and MV2 showed the highest diversity and richness (see also Table 2). Conversely, ACH and ERE samples showed lower diversity, reflecting a smaller range of viral taxa and potential dominance by a few species.

**Figure 3:**
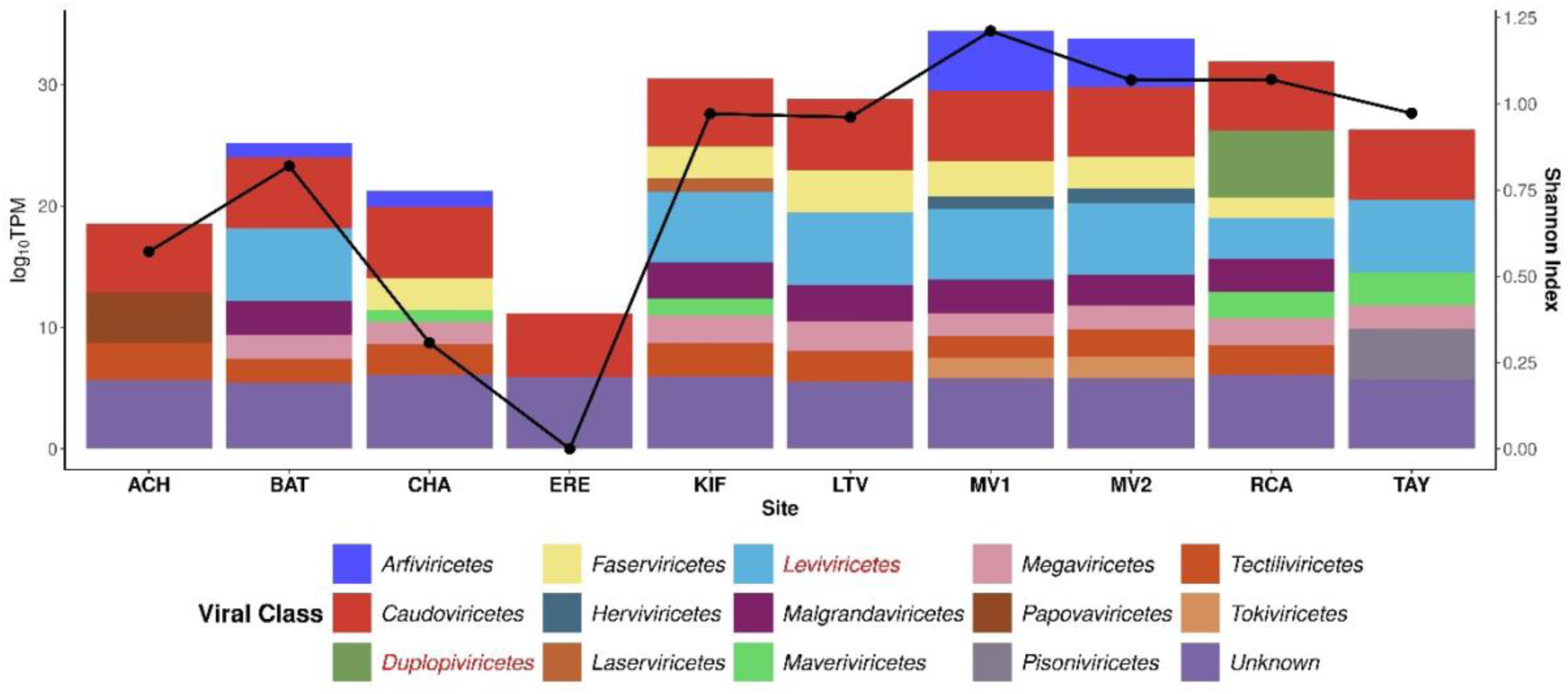
Stacked bar plot showing the taxonomic composition of mVBs at the class level across Antarctic soil samples (log scale), with Shannon α-diversity index overlaid as black dots. Note that a logarithmic scale was used to accommodate the large disparity between the “Unknown” viruses and the other classes, allowing patterns to be observed even when one group is very dominant. RNA viral classes are indicated in red and DNA classes in black in the legend.

To better understand how local factors contribute to shaping the structure and diversity of viral populations across the ice-free Antarctic soils, a t-SNE and PAM clustering (K=6) analysis of viral bin profiles were performed (Figure 4a). Six distinct clusters resolved using PAM clustering with K=6, achieving the best silhouette value. These clusters highlight the site-specific composition of viral communities, reflecting the ecological and environmental heterogeneity of Antarctic ice-free soils. ISA analysis showed in Figure 4b revealed that Cluster 1 (ACH, ERE) was characterized by *Caudoviricetes* viruses infecting primarily *Proteobacteria* and *Firmicutes*. Cluster 5 (LTV, RCA, TAY) also grouped *Caudoviricetes* associated with *Proteobacteria* and *Actinobacteriota*, although with lower diversity. In contrast, Cluster 6 (MV1, MV2) showed a broader viral and host diversity, combining multiple bacterial and archaeal phyla, and included RNA viral classes such as *Leviviricetes* and *Malgrandaviricetes* infecting *Proteobacteria*, *Actinobacteriota (Actinomycetota)*, and *Chlamydiota*. These patterns highlight how distinct viral groups and host affiliations underpin the ecological differentiation of Antarctic soil viral communities.

**Figure 4:**
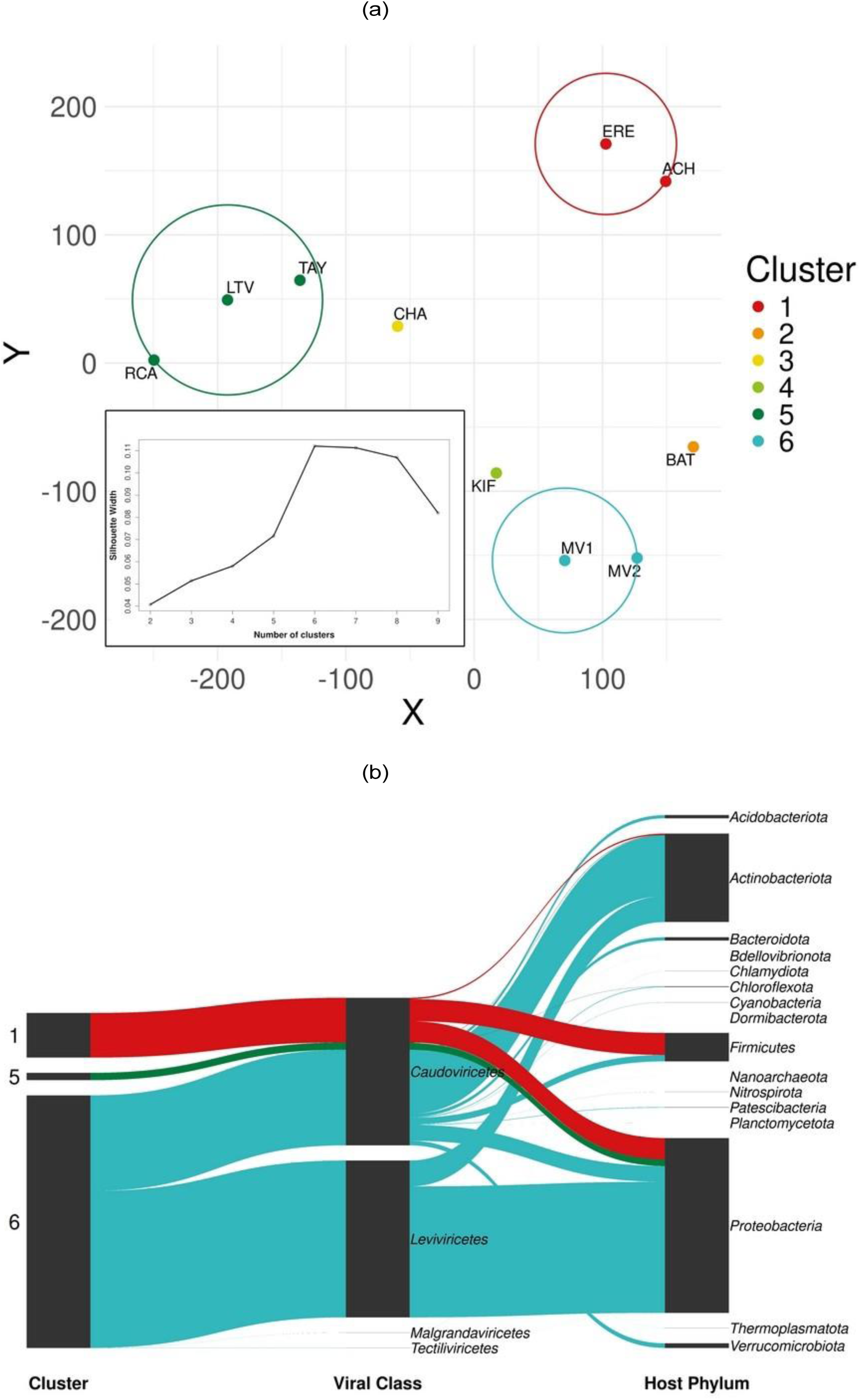
Clustering of viral bins from Antarctic soils. **(a)** t-SNE visualization with PAM clustering, based on Jaccard distances; the optimal number of clusters (k = 6) was selected based on the highest silhouette value, shown in the embedded plot at the bottom left. **(b)** ISA (p-value < 0.05; 1,000 permutations) results annotated with viral taxonomic classes and predicted host phyla, using the sum of TPM (Σ TPM) as a measure of relative abundance.

Host prediction analysis showed that most of the viruses have an unknown host and detected viruses infecting Bacteria, Archaea and eukaryotes, the most abundant being those infecting Bacteria (Figure S1 and S2). Among these, *Proteobacteria* was the most frequently annotated host phylum, closely followed by *Actinobacteriota*. The most abundant archaeal phylum was *Thermoproteota* and the only eukaryotic hosts predicted belonged to the phylum *Ochrophyta*.

### Environmental drivers of bins, AMGs and pathways distribution

To assess the influence of environmental factors on the mVBs, the AMGs, and their associated pathways, an RDA analysis was employed (Figure 5). The analysis revealed that the distribution of mVBs is significantly driven by elevation above sea level (p-value=0.03), explaining 60% of the observed variance (Figure 5a). This highlights the influence of altitude-related factors, such as temperature, on viral community composition. Sites at higher elevations are more likely to be subjected to harsher environmental conditions, leading to unique adaptations in the associated viral populations.

**Figure 5:**
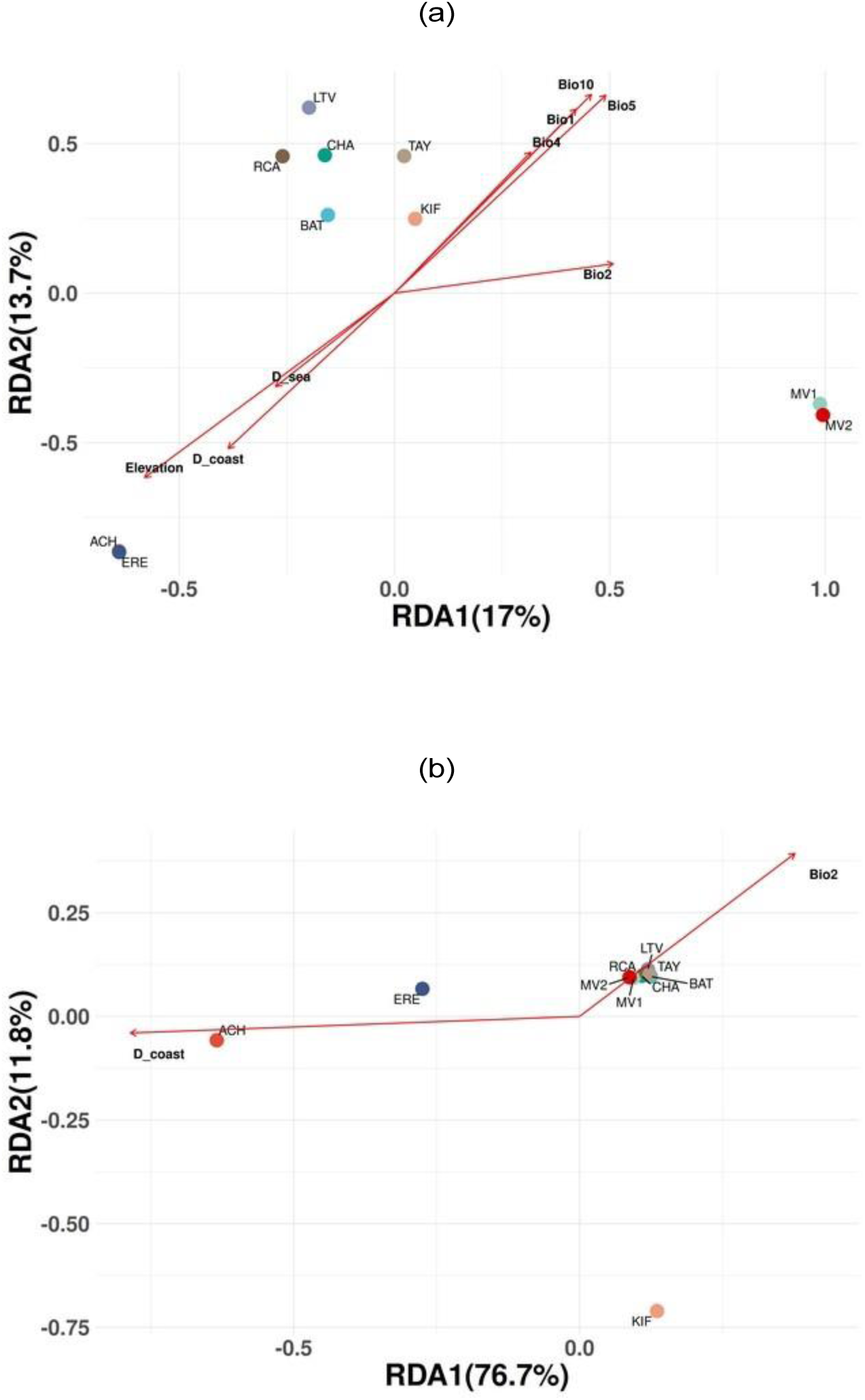
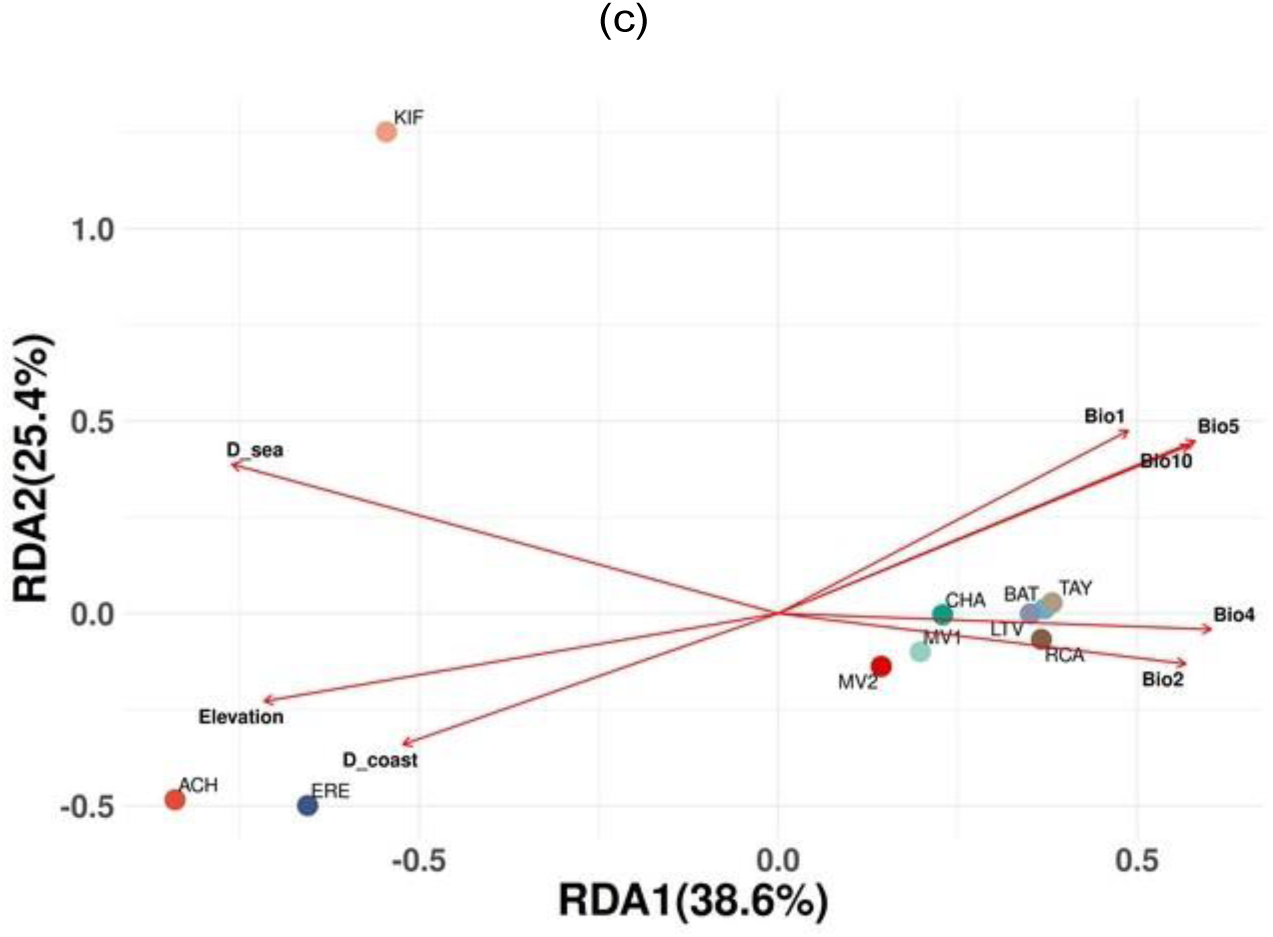
RDA analysis of **(a)** mVBs, **(b)** AMGs, and **(c)** metabolic pathways in Antarctic soils. Only environmental variables with a p-value ≤ 0.05 based on the permutation test (n = 1000) were included in the ordination plots.

For AMGs, distance to the coast was the most important variable (p-value=0.03), explaining 97% of the variance (Figure 5b), thus suggesting that proximity to the coastline—and associated gradients in salinity, moisture availability, and organic matter—strongly shapes the functional potential of viral AMGs in Antarctic soils. Similarly, RDA for metabolic pathways identified distance to the sea as the key driver (p-value=0.02), accounting for 84% of the variance (Figure 5c). These results emphasize how environmental gradients not only influence the taxonomic diversity of viruses but also determine their metabolic roles and contributions to ecosystem functioning.

### Host-Virus-AMG network analysis

A tripartite network analysis to link viral bins, AMGs and predicted host phyla was performed, comprising 9,732 nodes and 10,221 edges (Figure 6). This model provides an in-depth understanding of how viruses enhance host metabolic capabilities through their AMGs, revealing a complex interplay between viral diversity, host specificity, and functional potential.

**Figure 6:**
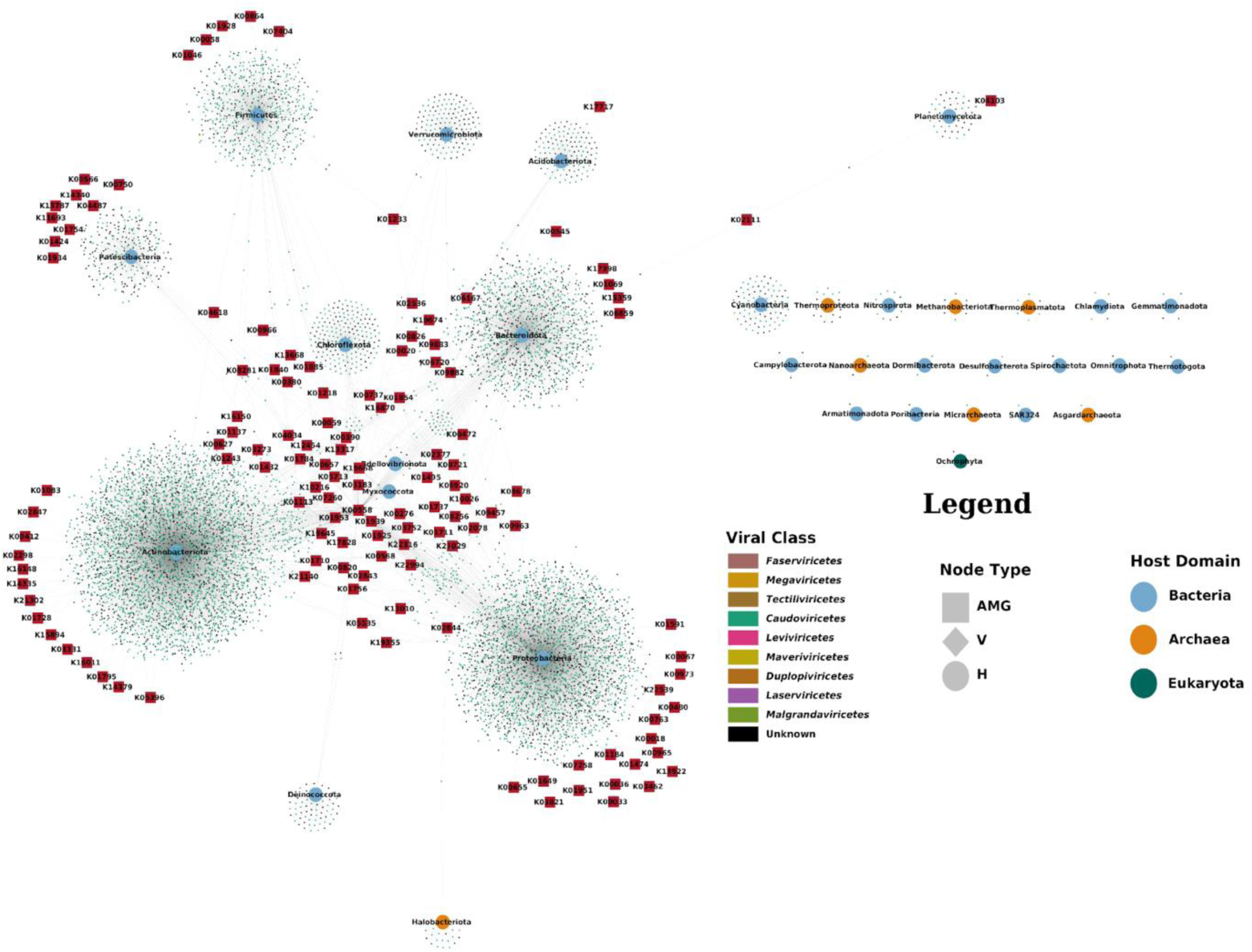
Network representing the relationship between mVBs and their detected AMGs and predicted host for all taxonomically annotated mVBs at host phylum level. V: virus; H: host.

The network showed several AMGs shared across multiple viruses infecting different host phyla, highlighting their ecological significance. Key hubs indicate highly interconnected AMGs central to critical metabolic pathways, such as cysteine and methionine metabolism. At the core of the network lies the *dcm* gene, encoding DNA (cytosine-5)-methyltransferase 1 (K00558; EC 2.1.1.37), which plays a pivotal role in cysteine and methionine metabolism (map00270). This gene is primarily associated with *Caudoviricetes* viruses infecting nine bacterial phyla, including the most abundant hosts, *Proteobacteria* and *Actinobacteriota*.

Peripheral regions of the network revealed AMGs unique to specific bacterial phyla, with *Proteobacteria* hosting the largest number of exclusive AMGs. In contrast, only one AMG was identified for an archaeal virus (*waaG*; K02844; EC 2.4.1.-), and none was found for eukaryotic viruses. Remarkably, none of the AMGs detected in this study were associated with RNA viral bins.

The network confirmed that bacterial phyla dominate as hosts, particularly *Proteobacteria* and *Actinobacteriota*, which are highly abundant in Antarctic soils. Archaeal hosts, such as *Thermoproteota*, were less frequently represented, and for eukaryotic hosts, only the phylum *Ochrophyta* was identified.

### The Antarctic Soil Auxiliarome

The complete catalogue of soil pan-AMGs from Antarctic ice-free areas, defined for the first time in this work as the auxiliarome, consisted of 2,851 AMGs coding for 219 different metabolic enzymes (Figure 7 and Table S6). This bipartite network (317 nodes; 360 edges) connects AMGs with their respective metabolic pathways, revealing diverse host metabolic engineering potential. Prominent pathways included 1) cysteine and methionine metabolism (map00270), driven by the *dcm*, and 2) nicotinate and nicotinamide metabolism (map00760), supported by *NAMPT* (K03462; EC 2.4.2.12; nicotinamide phosphoribosyltransferase) and *nadM* (K13522; EC 2.7.7.1; bifunctional NMN adenylyltransferase/nudix hydrolase) genes. The role of *ilvB* (K01652; EC 2.2.1.6; acetolactate synthase I/II/III large subunit), contributing to pantothenate and CoA biosynthesis (map00770), valine, leucine and isoleucine biosynthesis (map00290), butanoate metabolism (map00650) and C5-branched dibasic acid metabolism (map00660) is also quite relevant.

**Figure 7:**
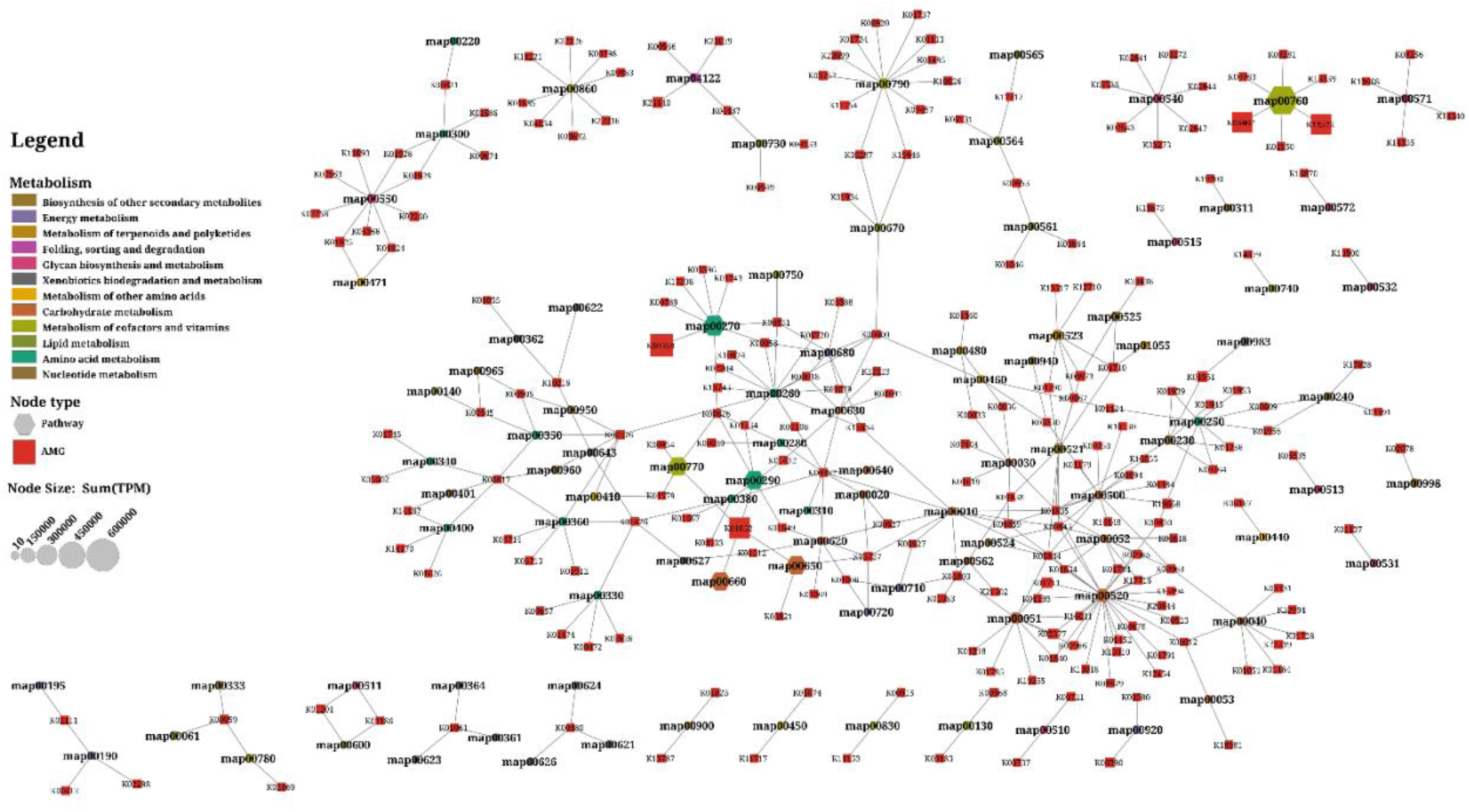
Auxiliarome network of Antarctic soils (pan-AMGs catalogue), showing the relationships between AMGs. AMGs are represented by KO identifiers, and metabolic pathways are represented by KEGG pathway names.

The entire metabolic potential of Antarctic soil AMGs was visualized in a global metabolic pathway map (Figure S3) that integrates all detected AMGs into broader metabolic contexts, providing insights on their roles in host adaptation and survival.

The t-SNE, PAM and ISA analysis of AMGs (Figure S4) reveals that viral auxiliary functions are organized according to environmental pressures and metabolic specialization. Cluster 1 (ACH, ERE) was dominated by AMGs associated with carbohydrate metabolism (e.g., K01711, *gmd*; K02377, *fcl*), amino acid metabolism (e.g., K01474, *hyuB*), and cofactor and vitamin biosynthesis (e.g., K03462, *NAMPT*; K13522, *nadM*), along with functions related to xenobiotic degradation (e.g., K00480, salicylate hydroxylase). In contrast, Cluster 2 (BAT, CHA, Mt. Kyffin (KIF), LTV, MV1, MV2, RCA and Taylor Valley (TAY)) was enriched in AMGs involved in lipid metabolism (e.g., K00059, *fabG*), energy metabolism (e.g., K00390, *cysH*), and cofactor and vitamin pathways (e.g., K01737, *queD*; K09882, *cobS*), suggesting viral adaptations that support host energy regulation and resilience under environmental stress. These patterns highlight how viral-mediated metabolic capacities align with distinct ecological strategies across Antarctic soil environments.

Similarly, t-SNE, PAM and ISA analysis of AMG-derived pathways (Figure S5) showed that Cluster 1 was dominated by pathways related to cofactor and vitamin metabolism (e.g., map00760, nicotinate and nicotinamide metabolism) with minor contributions from amino acid metabolism (e.g., map00471, D-glutamine and D-glutamate metabolism) and xenobiotic degradation (e.g., map00621, dioxin degradation). Cluster 2 was characterized by pathways associated with amino acid metabolism (e.g., map00270, cysteine and methionine metabolism), carbohydrate metabolism (e.g., map00052, galactose metabolism; map00500, starch and sucrose metabolism), energy metabolism (e.g., map00920, sulfur metabolism), and additional cofactor and vitamin pathways (e.g., map00780, biotin metabolism; map00790, folate biosynthesis; map00860, porphyrin metabolism). Cluster 3 (KIF), in turn, was specifically associated with amino acid metabolism (e.g., map00290, valine, leucine and isoleucine biosynthesis). These patterns reflect distinct metabolic strategies encoded by viral communities across Antarctic soils, with AMGs and their pathways divided into two major groups that likely reflect environmental pressures and metabolic specialization.

We analyzed the contributions of individual samples to AMGs and their associated metabolic pathways by examining both the number of AMGs detected (indicated by numbers adjacent to the bars in Figure 8) and their relative abundance (reflected by the bar lengths in Figure 8 and the color intensity in Figure 9). ACH and KIF samples showed significant contributions to the metabolism of cofactors and vitamins (e.g., *NAMPT*, *nadM*, and *ilvB*), amino acids (e.g., *dcm* and *ilvB*), and carbohydrates (*ilvB*); (see also Table S6). Cofactor and vitamin metabolism, with 426 AMGs (43 unique; Figure S6), was remarkable in terms of abundance, followed by carbohydrate metabolism with 526 AMGs (71 unique; Figure S7). Although amino acid metabolism contributes the largest number of AMGs with a total of 1,100 (55 unique; Figure S8), its overall abundance is comparatively lower.

**Figure 8:**
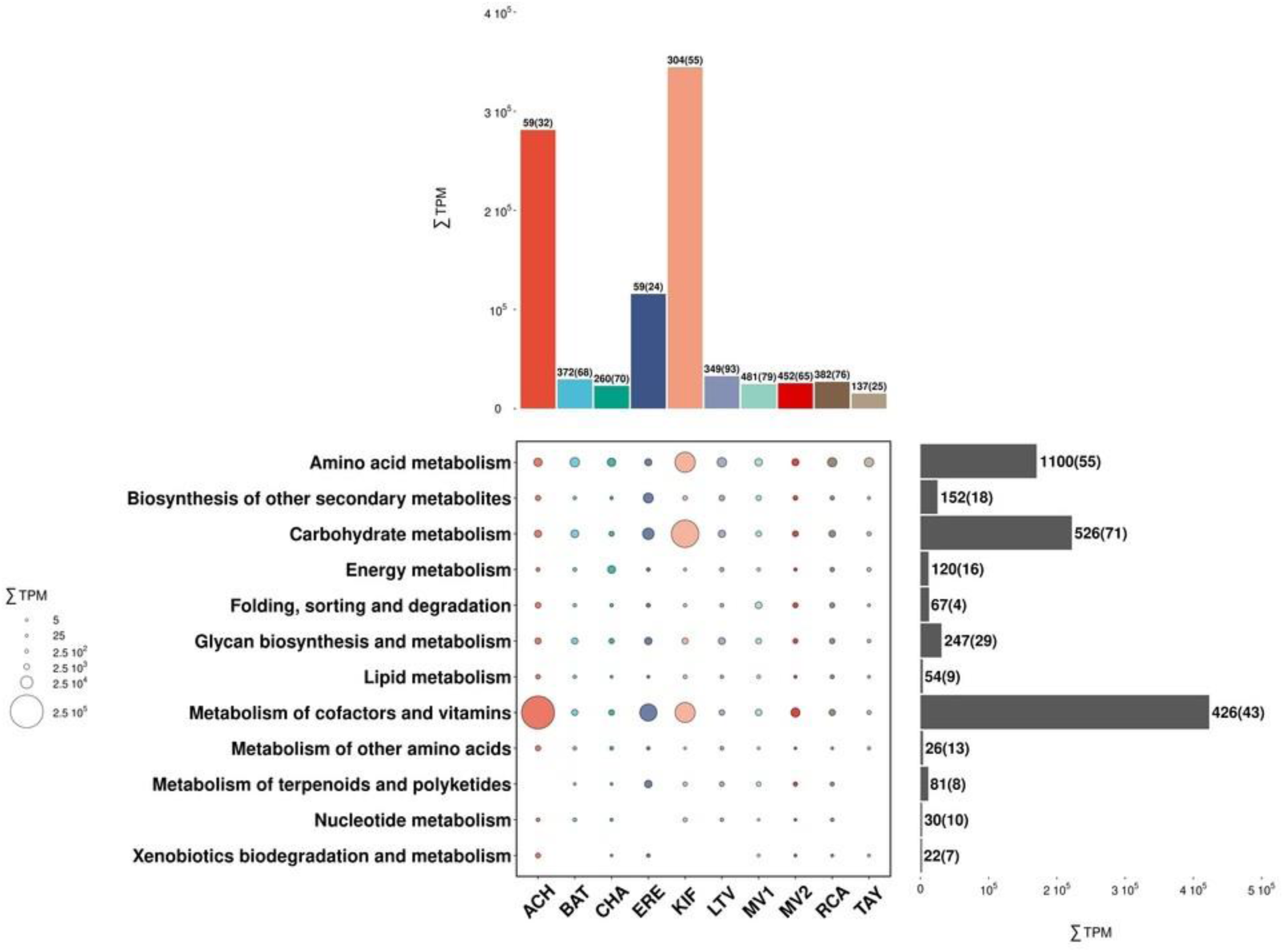
Bubble plot summarizing the contributions of Antarctic soil sampling sites to the metabolisms associated with AMGs, measured as relative abundances (ΣTPM). Numbers adjacent to the bars indicate the total number of genes, with the number of unique genes shown in parentheses.

**Figure 9:**
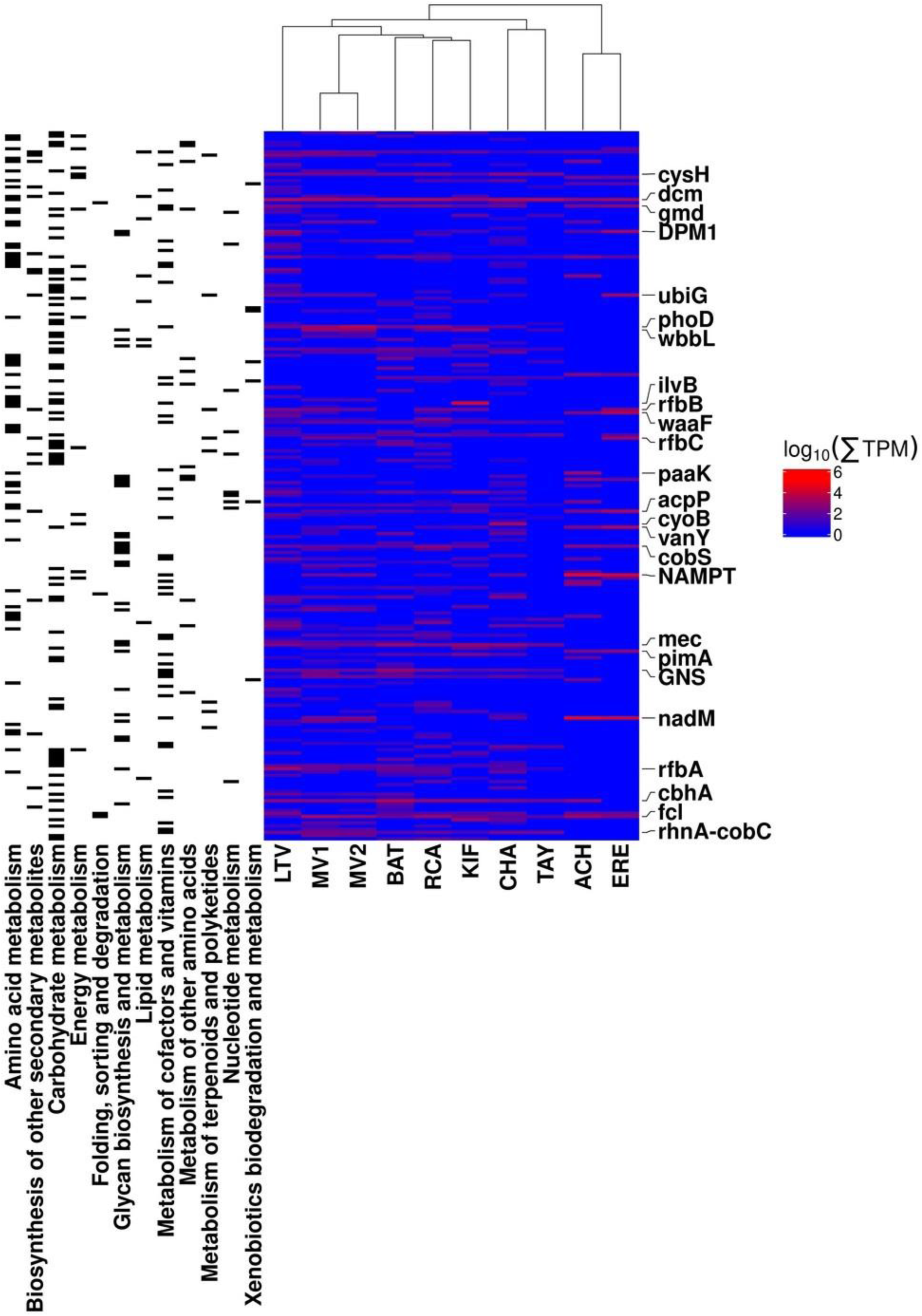
Heatmap showing the contributions of Antarctic soil sampling sites to AMGs, expressed as relative abundance (ΣTPM). Only the top 25 most contributing AMGs are labelled. Hierarchical clustering of sampling sites was performed using the complete linkage method and Jaccard distance.

A significant number of AMGs related to sulfur metabolism were detected (31 unique genes; Figure S9). Although sulfur metabolism does not correspond to a single standalone KEGG pathway category, it is distributed across several pathways involving the biosynthesis and degradation of sulfur-containing molecules, such as cysteine and methionine metabolism, folate biosynthesis, glutathione metabolism, sulfur metabolism, thiamine metabolism, and the sulfur relay system. The prominence of sulfur-associated AMGs highlights the central role of viral metabolic engineering in modulating sulfur cycling within Antarctic soil environments.

## Discussion

The samples analyzed in this study represent a unique transect of ice-free Antarctic soils, spanning elevations from sea level (Cape Hallet) to nearly 2,700 meters above sea level (Mt. Erebus) and geographical locations from coastal Oates Land to the Central TAM, passing through the arid McMurdo Dry Valleys. This range, to our knowledge, constitutes the most extensive study of the Antarctic soil virosphere published to date (see Supplementary Text, Comment 1).

### Taxonomic diversity of viral communities

The majority of bins identified in this study belong to unknown viral classes, highlighting the unique and largely unexplored nature of Antarctic soil viral communities. Among the identified DNA viruses, *Caudoviricetes* was the most frequently assigned class. All annotated RNA bins were classified under the classes *Leviviricetes* and *Duplopiviricetes*, which further emphasizes the taxonomic novelty of the viral populations uncovered.

A comparison with previous studies of Antarctic soil viral communities and other Antarctic ecosystems (Table S7) [9, 84–96] revealed that 13 of the 34 viral families identified in this study had been previously described in Antarctic soils.

In contrast, 18 families (highlighted in blue in the "Viral Family" column of Table S7) are reported here for the first time in Antarctica, and 3 more in Antarctic terrestrial environments (highlighted in red), expanding the known diversity of viruses in this extreme ecosystem. Additionally, a total of 14 other viral families previously reported in Antarctic soils or soil-related environments [9, 84, 93, 95] were not detected in this survey (see families highlighted in bold in Table S7, “Other Families” column).

The differences between the viral families identified in this work and those reported in previous studies of Antarctic soils could be attributed in part to the geographic and environmental heterogeneity of the selected sampling sites. This likely contributed to differences in the observed viral taxa. The inclusion of diverse ice-free soils, ranging from sea level to high elevations and spanning different moisture regimes, provided a broad representation of Antarctic viral communities not covered by earlier studies. Also, improvements in recent years in bioinformatic tools developed for identifying viral sequences could also be related to these observed differences. And we must not forget that the viral purification techniques employed––which may or may not include a pre-amplification step prior to sequencing––may also be related to these differences. Although we know that amplification can affect evenness [97], it is an effective strategy when dealing with low biomass environments and allows for the detection of greater viral diversity. In this case, we believe it is the best methodology to employ due to the oligotrophic conditions that characterize ice-free soils in Antarctica [98].

Table S7 shows that a greater number of viral families were detected in our study compared to previous studies conducted in Antarctica in soil and other environments. Also, this table shows that concentrating and amplifying viral particles yields higher diversity [9, 84–96], demonstrating the value of this methodology when the primary goal is to maximize taxonomic detection.

### Environmental factors driving viral distribution

Our analysis suggests that elevation above sea level is the main factor influencing the distribution of viruses in ice-free soils in Antarctica. This finding aligns with previous studies that have identified elevation as a critical determinant of microbial diversity in these regions [11]. Elevation is often correlated with changes in average temperature, UV radiation, and water availability, all of which are critical factors shaping viral community composition. Previous research has shown that calcium content and pH also play significant roles in determining the distribution of viral communities in Antarctic soils [4].

Interestingly, elevation has been identified as the primary driver of bacterial distribution in Antarctic soils [6]. This pattern is consistent with our findings, as Bacteria are the main hosts for the viruses detected. Similarly, in the Taylor Valley—located at the McMurdo Dry Valleys— elevation was found to influence both bacterial and fungal communities [99]. For Archaea, which represent the second most frequently detected viral host group, diversity was positively correlated with soil water content in McMurdo soils [100]. This result is consistent with our findings, as elevation is moderately correlated with soil water content in the TAM region [101]. Protist diversity, in contrast, has been shown to decrease with elevation in McMurdo Dry Valleys soils [102], further highlighting the complex interplay between elevation and microbial community composition.

In contrast, no decrease in viral diversity was detected with increasing southern latitude, highlighting the counterintuitive notion that latitude is not a primary driver of viral diversity in ice-free Antarctic soils. A similar pattern was previously observed in freshwater Antarctic lake viromes [89].

Unlike elevation, distance to the coast was the main factor explaining the distribution of AMGs across samples. However, in Antarctica, the presence of extensive sea ice shelves complicates the interpretation of this variable. For instance, although the ACH and KIF samples are relatively close to the Ross Sea shoreline (123 km and 8 km, respectively), they are separated from the open sea by 680 km and 636 km due to the Ross Ice Shelf. Thus, distance to the sea—rather than distance to the coast—emerges as the most relevant factor in explaining AMGs and metabolic pathway distributions. Both variables, however, likely correlate with soil moisture content, salinity, and temperature gradients, which may influence viral metabolic engineering strategies.

The t-SNE analysis of AMGs emphasizes the role of viral AMGs in shaping host metabolic processes across Antarctic ice-free soils, with clear spatial and functional segregation in extreme environments. This study also demonstrates that clustering patterns observed in the t-SNE and PAM analyses are consistent with those from RDA for mVBs, AMGs, and metabolic pathways. For example, the clustering of Miers Valley samples (MV1 and MV2) reflects their shared sampling location and environmental conditions. Similarly, ERE and ACH samples cluster together due to their high elevation, low α-diversity, lack of RNA mVBs, and shared trends in temperature and precipitation seasonality (bio1, 2, 5, 10, 15). In contrast, KIF sample stands out as an outlier, likely due to the high proportion of endemic mVBs, particularly among the RNA bins.

### Functional insights from AMGs

The analysis of AMGs in Antarctic ice-free soils revealed several genes with critical roles in host metabolism, highlighting their importance in viral-mediated ecological processes. Among the most notable AMGs are *dcm*, *NAMPT*, *nadM*, and *ilvB*, each with distinct metabolic contributions and ecological and functional implications:

- *dcm*: This gene, present at all sampling sites, is predominantly found in *Caudoviricetes* infecting 11 predicted bacterial phyla. Dcm catalyzes the transfer of a methyl group from S-adenosyl-L-methionine to cytosine, forming S-adenosyl-L-homocysteine and methylated DNA. This reaction is part of cysteine and methionine metabolism and also plays and important role in DNA methylation as part of restriction-modification systems that protect bacteria against bacteriophage infections. Previously described roles for bacteriophage-encoded methyltransferases include: (1) methylation of viral genomes to evade host restriction enzymes, (2) modulation of host epigenetic landscapes, and (3) regulation of viral genome replication and encapsidation [103–107]. Beyond DNA regulation, viruses may redirect organic sulfur from methionine into cysteine—which is the only amino acid that can form disulfide bonds—helping assemble cysteine-rich structural proteins that stabilize virions in energy-limited anoxic habitats (Ashcroft et al., 2005). This gene likely provides a fitness advantage to viral populations (see Supplementary Text, Comment 2). Sánchez-Romero et al. [106] determined that it is present in one fifth of phage genomes.
- *NAMPT* and *nadM*: The enzymes encoded by these genes are central to the salvage biosynthetic NAD^+^ pathway, which is crucial for cellular REDOX reactions and as a cofactor for numerous enzymes. NAMPT catalyzes the transformation of nicotinamide D-ribonucleotide and diphosphate into nicotinamide mononucleotide (NMN) and 5-phospho-alpha-D-ribose 1-diphosphate. NadM converts NMN and ATP into NAD^+^ and diphosphate. These AMGs were primarily detected in *Caudoviricetes* infecting *Proteobacteria*, suggesting that enhancing NAD^+^ biosynthesis might benefit Antarctic soil viruses by supporting host metabolism under nutrient-limited conditions.
- *ilvB*: This gene encodes an enzyme involved in the biosynthesis of branched-chain amino acids (valine, leucine, isoleucine), carbohydrate metabolism, and pantothenate/CoA biosynthesis. It was only detected in the KIF sample, in an abundant unclassified mVB with unknown host. It catalyzes the decarboxylation of pyruvate to form acetolactate and CO_2_, an essential step in multiple pathways. The restricted presence of *ilvB* in KIF suggests that modulation of these functions may confer significant metabolic advantages to viruses in this region, potentially influencing host energy metabolism, stress resistance, and cellular regulation.

Other interesting AMGs worth mentioning include the gene encoding cellulose 1,4-beta-cellobiosidase (*cbhA*), involved in starch and sucrose metabolism. This enzyme plays a crucial role in the breakdown of the non-reducing ends of cellulose to release cellobiose, which can then be hydrolyzed into glucose, an important source of energy for the cell. CbhA also has biotechnological implications, as it catalyses one of the limiting steps in biofuel production using plant-derived byproducts [108]. It was detected mainly in *Caudoviricetes* infecting *Actinobacteriota* and *Proteobacteria*. The presence of *cbhA* in Antarctic soil viruses suggests a viral contribution to facilitating energy acquisition from cellulose-degrading bacteria.

Also remarkable is the *vanY* AMG. This AMG, detected in *Caudoviricetes* infecting Bacteria (e.g., *Actinobacteriota*, *Bacteroidota*, *Firmicutes*, *Patescibacteria*, and *Proteobacteria*), is associated with peptidoglycan biosynthesis and vancomycin resistance. VanY modifies the D-Ala-D-Ala termini of peptidoglycan precursors, thereby preventing vancomycin binding and allowing bacterial cell wall cross-linking to proceed [109]. Since vancomycin is primarily produced by *Actinobacteriota*, especially *Actinomycetes* [110], viral modulation of this function may represent a strategic adaptation to help hosts resist antibiotic-producing competitors. Furthermore, our findings suggest that vancomycin resistance mediated by *vanY* may extend to previously unrecognized bacterial phyla [111], underscoring the ecological relevance of AMGs in mediating microbial interactions and resource competition in Antarctic soils.

We also identified several AMGs involved in antibiotic production (see Supplementary Text, Comment 3) and in organic and inorganic sulfur metabolism (Table S6 and Figure S9) (see Supplementary Text, Comment 4). Sulfur-related AMGs may play crucial roles in the biogeochemical sulfur cycle and may enhance the adaptability of viral hosts to nutrient-poor Antarctic soils.

All these findings underscore the critical role of viral AMGs in driving host metabolic engineering and maintaining ecosystem functionality under the extreme conditions of Antarctic ice-free soils. Interestingly, no AMGs were identified in RNA viral bins.

### Diversity of predicted viral hosts

The predicted viral hosts include a wide range of bacterial and archaeal phyla, highlighting the extensive host range of Antarctic soil viruses. Nearly all of the predicted bacterial phyla had been described in previous research [6], with the exception of *Omnitrophota*, *Poribacteria*, and *Thermotogota*, emphasizing the novelty of some host-virus interactions in Antarctic soils (see Supplementary Text, Comment 5).

Among archaeal hosts, six phyla were predicted. Four of these—*Halobacteriota*, *Methanobacteriota*, *Thermoplasmatota*, and *Thermoproteota*—had previously been reported in the McMurdo Dry Valleys [100, 112, 113]. Notably, two archaeal phyla, *Asgardarchaeota* and *Micrarchaeota*, were identified here for the first time. *Asgardarchaeota* was predicted in a mVB assigned to *Caudoviricetes* from the LTV sampling site. *Micrarchaeota* was similarly predicted in a *Caudoviricetes* mVB from the KIF sample (see Supplementary Text, Comment 6).

Eukaryotic hosts were minimally represented in the dataset, with only a single unclassified virus infecting *Ochrophyta* detected at the LTV sampling site. *Ochrophyta* are photosynthetic algae previously reported in ice-covered lakes of the McMurdo Dry Valleys [114, 115] and in Antarctic soils [102, 116]. The detection of viral interactions with these algae suggests a potential ecological role for viruses in regulating eukaryotic microbial populations within Antarctic ecosystems. But, even when host microorganisms could not be predicted, several eukaryotic viruses were identified in our samples, including members of the families *Circoviridae*, *Herpesviridae*, *Iridoviridae*, *Mimiviridae*, *Nanoviridae*, *Papillomaviridae*, *Partitiviridae*, and *Phycodnaviridae*. In any case, compared to the number of prokaryotic viruses identified in this study, the frequency of eukaryotic viruses detected in our samples was very low, which is consistent with previous studies [9, 84, 88, 93, 95].

## Conclusions

This study highlights the critical role of viruses in regulating microbial communities across diverse Antarctic terrestrial regions. By covering a wide latitudinal transect, it represents the most extensive description of soil DNA and RNA viruses in Antarctica to date. We identified key environmental factors influencing viral distribution: (a) elevation above sea level as the primary driver of soil viral distribution, (b) distance to the coast as a likely determinant of AMG distribution among samples, and (c) distance to the sea as the main factor shaping the distribution of metabolic pathways. Additionally, we expand the known Antarctic soil virosphere by reporting the presence of 18 previously undescribed viral families in Antarctic soils.

Despite the harsh environmental conditions that characterize terrestrial environments in Antarctica—including low temperatures, freeze-thaw cycles, desiccation, nutrient limitation, and high UV radiation—our results indicate that viral communities in these ecosystems exhibit remarkable richness and functional potential. However, our understanding of the full extent of this viral diversity remains incomplete, as highlighted by the high proportion of unclassified viruses identified in this study.

This work proposes the formulation of the “auxiliarome” concept, envisioned as a community-scale pan-AMG catalogue. This novel framework integrates the biochemical potential of viruses to modulate host metabolism and contribute to ecosystem-level processes. The Antarctic ice-free soil auxiliarome demonstrates the vast potential of viruses to engineer host metabolism in extreme environments. This includes contributions to the metabolism of cofactors and vitamins, amino acids, carbohydrates, and sulfur, as well as adaptive traits beyond traditional metabolic pathways, such as defense against restriction-modification systems and antibiotic resistance. These functions are essential to microbial activity, diversity, and ecosystem stability.

## Acknowledgements

We would like to thank Prof Hugh Morgan and Prof Ian Hogg (Waikato University) for collecting samples at the Central TAM.

We thank the staff of the Electron Microscopy Service at Centro de Biología Molecular Severo Ochoa (CBM, CSIC), Madrid for the preparation of the TEM samples, the Center of Scientific Computation of the Universidad Autónoma de Madrid for hosting the HPC cluster employed in this work and the Biocomputational Analysis Facility at CBM for upload the reads to the ENA (European Nucleotide Archive). We thank Josefa Anton, from the University of Alicante, for her support in this project.

S.S.C. was supported by a FPU (FPU18/04695) predoctoral grant awarded by the Spanish Ministry of Science, Innovation and Universities. R.G.S. was supported by the postdoc fellowship “Margarita Salas” (MARSALAS22) granted by the University of Alicante. Financial support was provided by grant 24-UOW-033 from the New Zealand Marsden Fund to I.R.M. and A.A.

## Data availability

All the raw reads can be found at the European Nucleotide Archive under the study accession identification PRJEB76654.

## Author contributions

Conceptualization, S.C.C. and A.A.; Funding acquisition, A.A., I.R.M., S.S.C. and R.G.S.; Sampling, S.C.C. and I.R.M.; Sample processing, R.G.S. and M.C.F.M.; Data analysis, S.S.C.; Supervision, A.A., I.R.M. and R.G.S.; Writing—original draft, S.S.C. and R.G.S.; Writing—review and editing, A.A., I.R.M., R.G.S. and S.S.C.

