## Supplemental Figure 1 for "Antarctic Soil Auxiliarome: unraveling the pan-auxiliary metabolic genes catalogue in a transect across different ice-free regions of Antarctica"

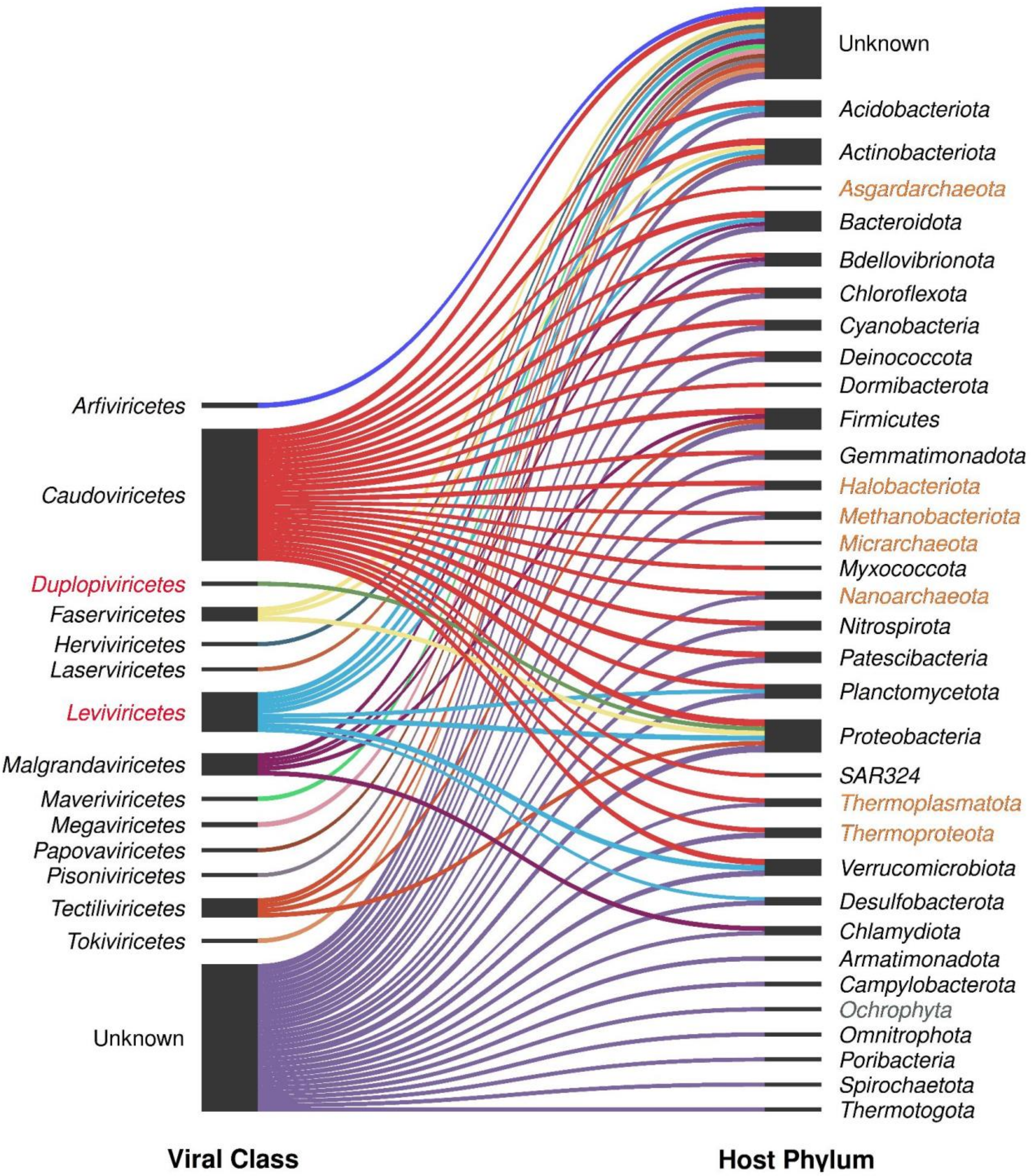


**Figure S1**: Sankey plot representing the relationship of mVBs and their predicted hosts. The width of the nodes represents the relative abundance on a logarithmic scale.
