## Supplemental Figure 2 for "Antarctic Soil Auxiliarome: unraveling the pan-auxiliary metabolic genes catalogue in a transect across different ice-free regions of Antarctica"

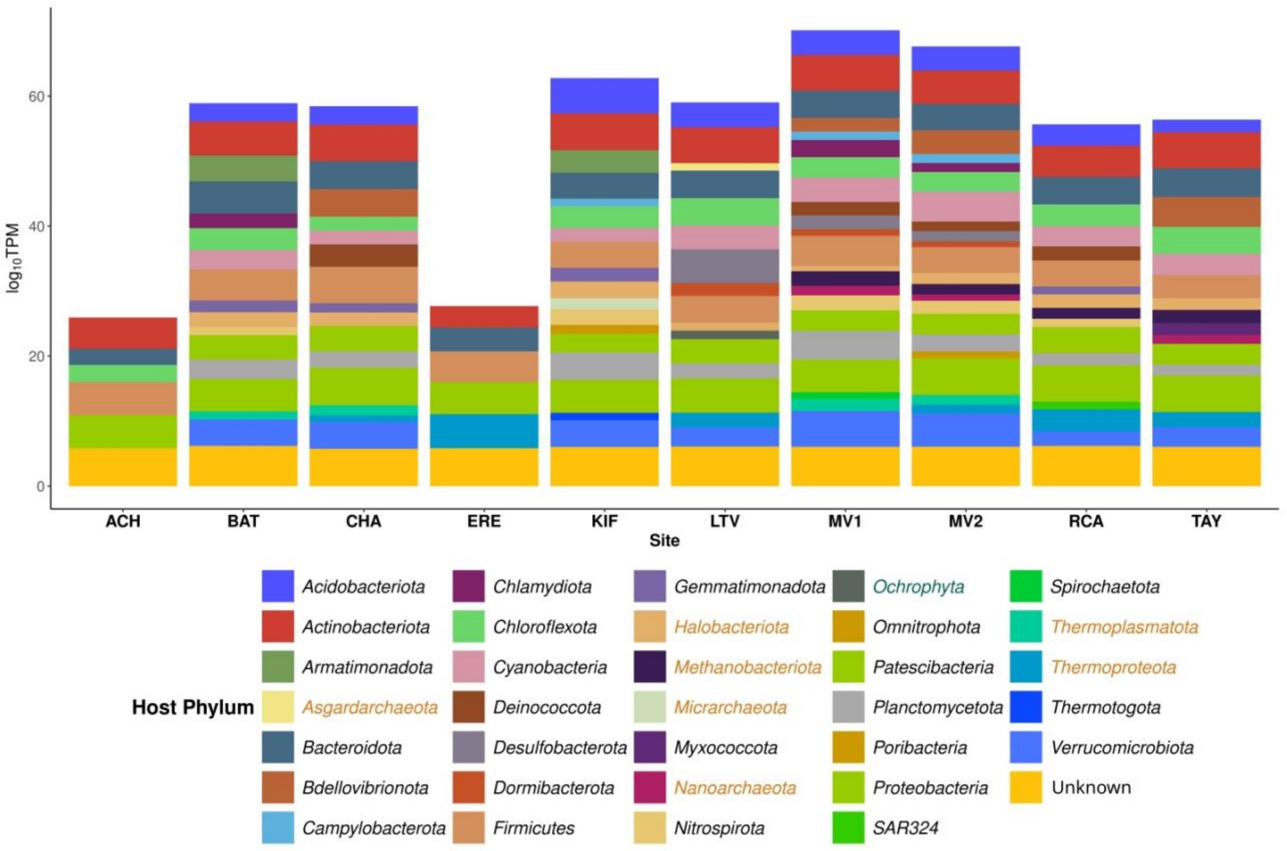


**Figure S2**: Stacked bar plot showing the predicted taxonomy of the host of the mVBs at the *Phylum* level on a logarithmic scale. The names in black represent *Bacteria*, the orange ones *Archaea* and the green one *Eukarya*.
