## Supplemental Figure 3 for "Antarctic Soil Auxiliarome: unraveling the pan-auxiliary metabolic genes catalogue in a transect across different ice-free regions of Antarctica"

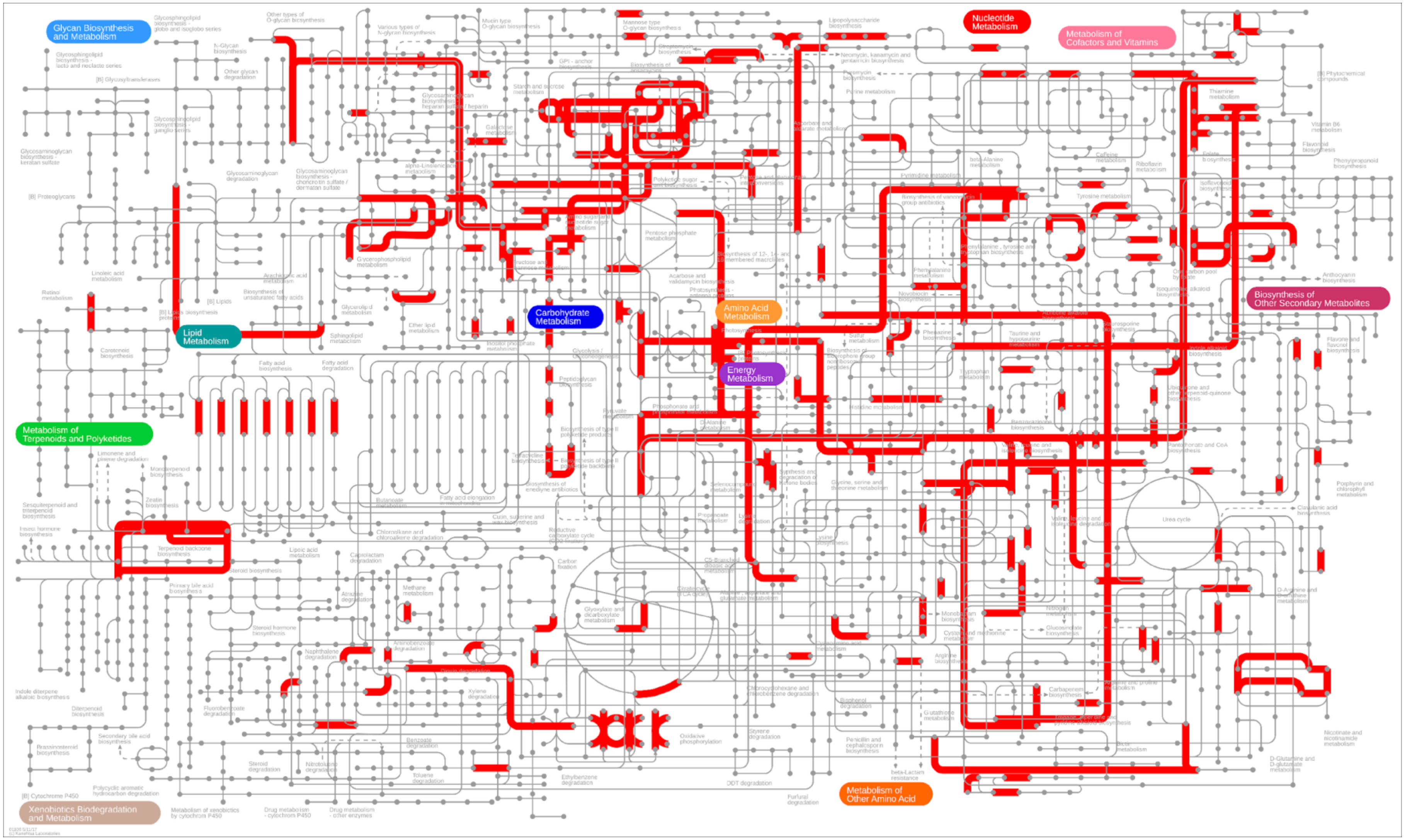


**Figure S3**: Metabolic map including the AMGs detected in this study. The red lines represent the segments of the metabolic pathways to which the AMGs belong.
