## Supplemental Figure 4 for "Antarctic Soil Auxiliarome: unraveling the pan-auxiliary metabolic genes catalogue in a transect across different ice-free regions of Antarctica"

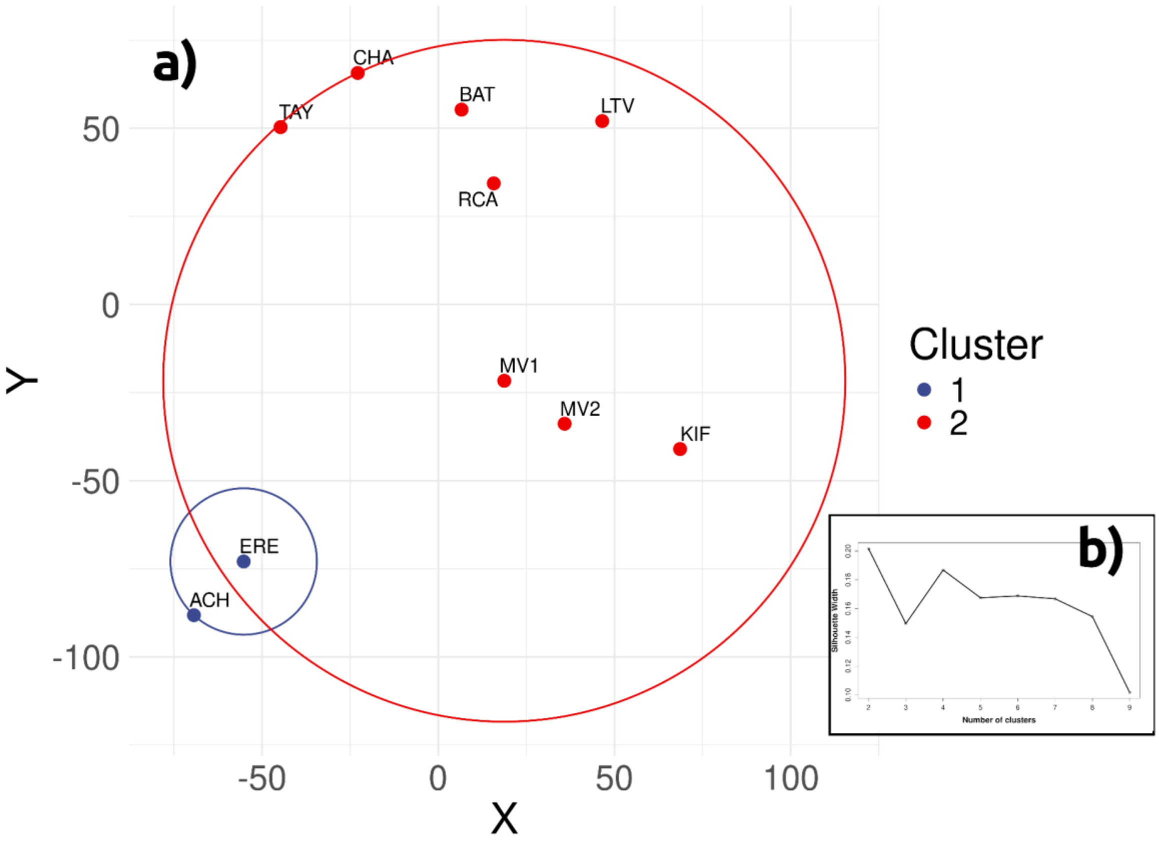


**Figure S4**: a) t-SNE analysis of AMGs with PAM clusters for K=2, obtained with the highest silhouette value, shown in b).
