## Supplemental Figure 5 for "Antarctic Soil Auxiliarome: unraveling the pan-auxiliary metabolic genes catalogue in a transect across different ice-free regions of Antarctica"

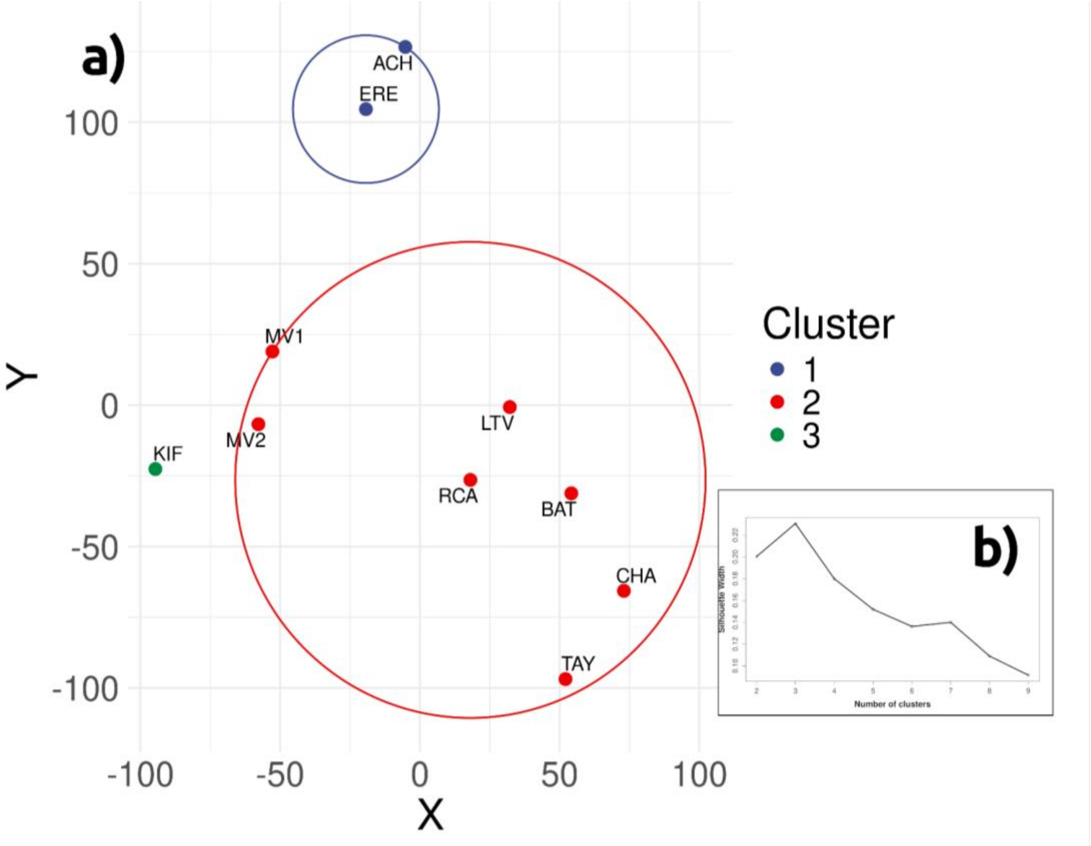


**Figure S5:** a) t-SNE analysis of AMG-derived pathways with PAM clusters for K=3, obtained with the highest silhouette value, shown in b).
