## Supplementary figures and images for "Antarctic Soil Auxiliarome: unraveling the pan-auxiliary metabolic genes catalogue in a transect across different ice-free regions of Antarctica"

### Supplemental Figure 6

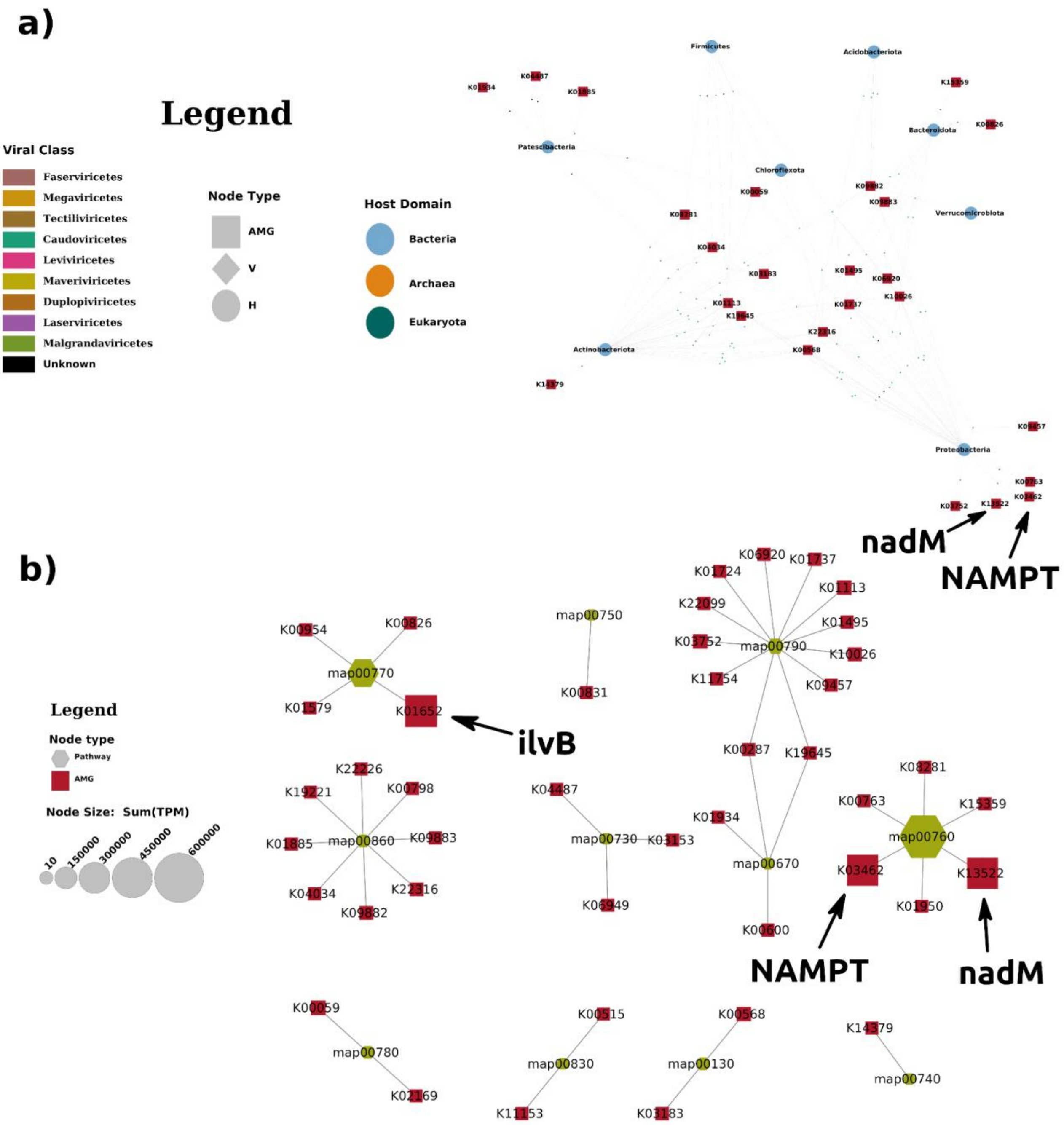


**Figure S6**: Metabolism of cofactors and vitamins. a) AMG-mVB-Host network and b) auxiliarome network.

### Supplemental Figure 7

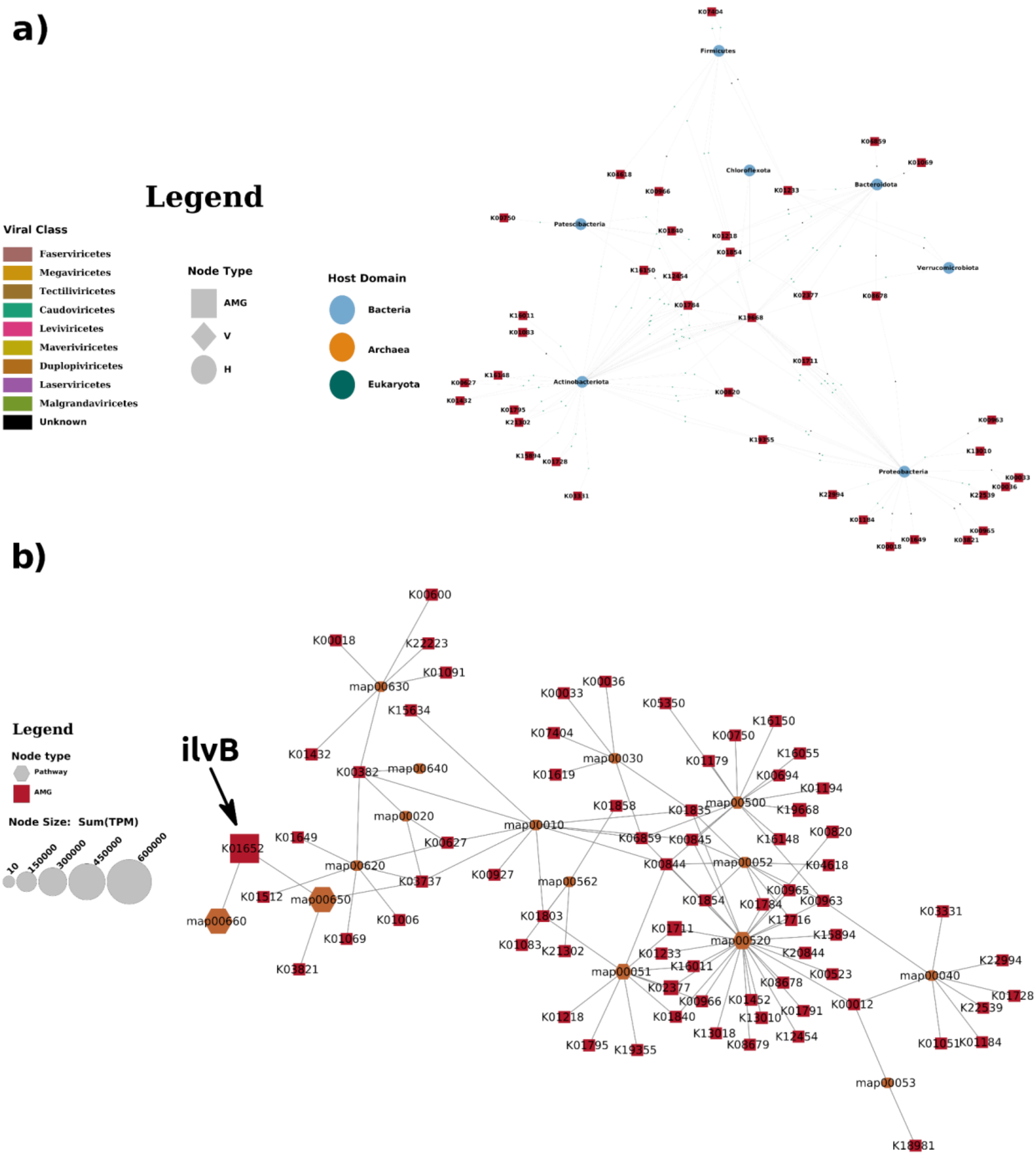


**Figure S7**: Metabolism of carbohydrates. a) AMG-mVB-Host network and b) auxiliarome network.

### Supplemental Figure 8

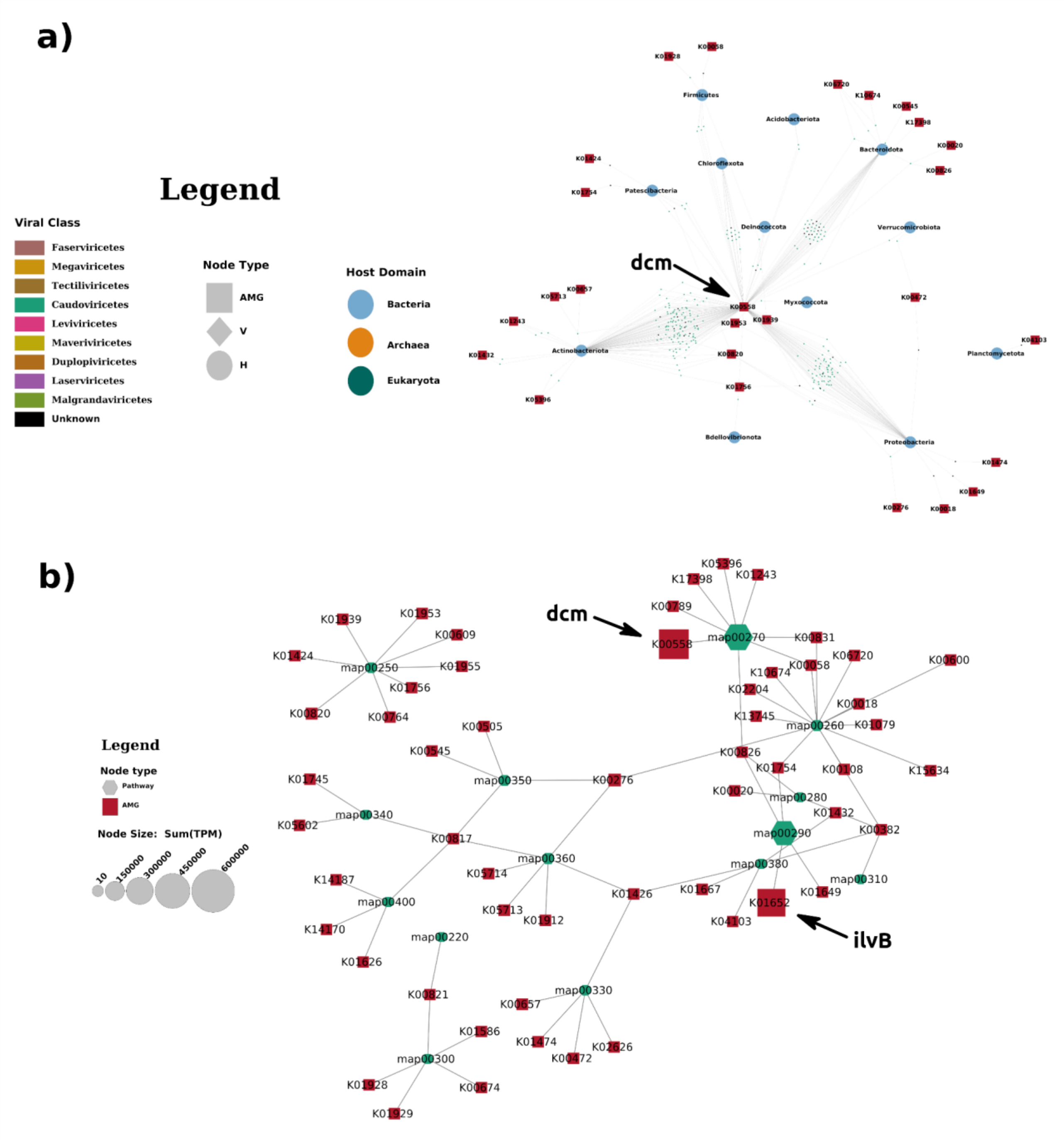


**Figure S8**: Metabolism of amino acids. a) AMG-mVB-Host network and b) auxiliarome network.

### Supplemental Figure 9

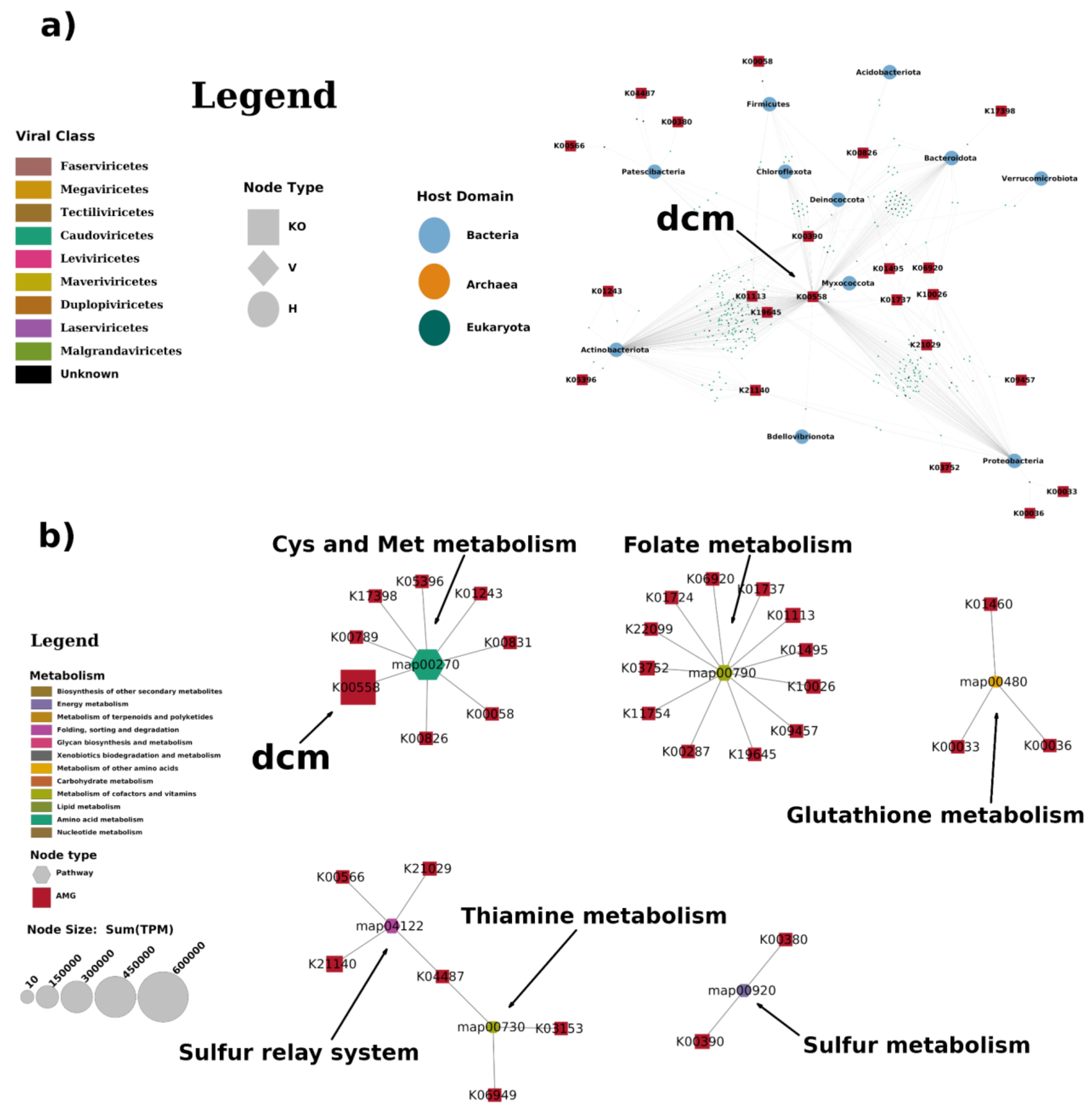


**Figure S9**: Sulphur related metabolism. a) AMG-mVB-Host network and b) auxiliarome network.
