## Supplemental Text for "Antarctic Soil Auxiliarome: unraveling the pan-auxiliary metabolic genes catalogue in a transect across different ice-free regions of Antarctica"

#### **Supplementary text**

### Comment 1

Ettinger et al. [1] studied endolith communities from the McMurdo Dry Valleys, Southern Victoria Land and Northern Victoria Land; Adriaenssens et al. [2] soil samples from the McMurdo Dry Valleys, Wei et al. [3] soils, hypoliths, chasmoendoliths and cryptoendoliths from the McMurdo Dry Valleys, while Zablocki et al. [4] studied hypoliths and surrounding soils from just one of the McMurdo Dry Valleys (Miers Valley). Therefore, our transect encompasses the most extensive study of the Antarctic soil virosphere published to date, due to the geographical extent of our sampling sites.

### Comment 2

Bacteria express sequence-specific endonucleases (REases) and the DNA methyltransferases that methylate identical sequences (MTases) at the same time, in such a way that bacterial methylated DNA escapes from digestion while exogenous viral DNA does not, constituting what is known as Restriction-Modification (R-M) Systems.

We have detected this enzyme in 814 abundant mVBs in a total of 51,559 mVBs (1.6%). According to Murphy et al. [5], one fifth of phage genomes include a MTase, so our detection rate is well below this figure. This discordance can be explained by the low number of completed genomes found in our work. Further analysis must be done to better understand the mechanisms and ecological implications behind the role of the MTases encoded by viruses in Antarctic soils.

### Comment 3

Among the AMGs involved in antibiotic production we can highlight the following:

a) Streptomycin (map00523; K01710: rfbB (28), K00973: rfbA (8), K00067: rfbD (7), K01790: rfbC (3), K00844: HK (1), K00845: glk (1), K01835: pgm (1) and K01858: INO1 (1)). We have detected these AMGs in *Caudoviricetes* with known hosts among *Actinobacteriota* and *Proteobacteria*. Streptomycin is produced by members of *Actinobacteriota* [6].

b) Validamycin (map00525; K01710: rfbB (28), K00973: rfbA (8) and K20438: valG (1)). This antibiotic in known to be produced by *Actinobacteriota* [7] and we have detected these AMGs in *Caudoviricetes* infecting *Actinobacteriota* and *Proteobacteria.*

c) Vancomycin (map01055; K01710: rfbB (28)). We have detected this AMG in *Caudoviricetes* infecting *Actinobacteriota* and *Proteobacteria*, however it was described in *Actinobacteriota* [8].

d) Aurachin (map00998; K02078: acpP (19)). This biomolecule is a a farnesylated quinolone alkaloid which is an excellent inhibitor of the respiratory chain with actibacterial and also antiprotozoal functions, whose described producers are members of *Myxococcota* and *Actinobacteriota* [9]. However, we have found it in viruses that infect *Proteobacteria* and *Verrucomicrobiota.*

e) Neomycin, kanamycin and gentamicin (map00524; K00844: HK (1) and K00845: glk (1)). These AMGs involved in the production of these aminoglycoside antibiotics have been found in unclassified viruses with unknown host, but the classical producers are within the *Actinobacteriota* phylum [10].

f) Novobiocin (map00401; K00817: hisC (1) and K14187: tyrA (1)). We found *Caudoviricetes* as a possible carrier with a non-detected host. This antibiotic is produced by members of *Actinobacteriota* [11].

g) Penicillin and cephalosporin (map00311; K19200: IAL (1)), detected in *Caudoviricetes* with unknown host and producer described within the fungi.

Our results suggest that viruses and AMGs may be relevant in the production of antibiotics in Antarctic soils and point to the presence of other unknown antibiotic producers in taxa other than those described above.

#### Comment 4

In this study we have found several AMGs regarding organic and inorganic sulphur metabolism. Specifically to cysteine and methionine metabolism: 948 (8 enzymes: K00558: dcm (926), K05396: dcyD (8), K17398: DNMT3A (4), K01243: mtnN (5), K00789: metK (2), K00826: ilvE (1), K00058: serA (1) and K00831: serC (1)), folate biosynthesis: 176 (12 enzymes: K01113: phoD (65), K10026: queE (36), K01737: queD (27), K06920: queC (14), K00287: folA (10), K11754: folC (5), K22099: folC2 (5), K19645: dfrB (1), K01495: folE (12), K03752: mobA (1), K01724: phhB (2), K09457: queF (1)), sulfur metabolism: 89 (2 enzymes: K00390: cysH (88), K00380: cysJ (1)), sulfur relay system: 67 (4: K21140: mec (48), K21029: moeB (15), K04487: iscS (3), K00566: trmU (1)), thiamine metabolism: 6 (3 enzymes: K04487: iscS (3), K06949: rsgA (2), K03153: thiO (1)) and glutathione metabolism: 3 (2: K00036: G6PD (1), K01460: gsp (1) and K00033: PGD (1)).

The main contributor was dcm, accounting for 926 appearances, followed by cysH with 86. Kieft et al. [12] found 227 AMGs (5: dsrA, dsrC/tusE, soxC, soxD and soxYZ) belonging to oxidation of inorganic sulfur and thiosulfate in samples distributed across different biomes, mainly in the ocean, and no one in polar or alpine soils. We have not found any AMG in common (adding up cysH and cysJ, which are involved in assimilatory sulfate reduction), that corresponds to the set of AMGs described by Kieft et al. [12]. Our results suggest that the pull of AMGs involved in the inorganic sulfur metabolism in Antarctic soils differs from the studied by Kieft et al. [12], probably due to the specificity of the Antarctic soil environmental constrictions.

#### Comment 5

*Omnitrophota* has been recognized in recent times as mainly inhabiting oligotrophic groundwater environments and sediments, but also present in soils, lakes, rivers, marine environments and even in wastewater [13]. It has been detected once in an unclassified mVB in KIF sample. Further research is needed to shed light on this phenomenon. *Poribacteria* was discovered by Fieseler et al. [14] in marine sponges and later other studies have detected them also in corals [15, 16]. In our samples it has been found in MV2 once, in an unclassified mVB. This sample is 9.5 km from the coast and 18.8 km from the sea, so wind transportation from marine aerosols could be a possible explanation, also supported by Chong et al. [17]. *Thermotogota,* a phylum definitively established by Oren et al. [18] based on Reysenbach et al. [19], is a phylum of thermophilic *Bacteria* and was also detected in only one unclassified mVB in KIF sample*,* where no geothermal activity has been described. These previously undetected phyla could be an error in the host prediction or a misclassification, for example with other members of the superphylum *Planctomycetota*-*Verrucomicrobiota*-*Chlamydiota* (PVC), to which *Omnitrophota* and *Poribacteria* also belong and with which they are closely related. A new discovery in Antarctic soils could also be a possibility, but further research must be done in order to shed light on this issue.

#### Comment 6

*Asgardarchaeota* members have been found in marine and marine-derived habitats, mangrove sediments, saline microbial mats and sediments, lake sediments, hydrothermal vents and cold seeps [20]. However, *Micrarchaeota* have been described in geothermal sites, soils, hypersaline environments and freshwater and are known to be metabolically dependent and physically associated with member of the phylum *Thermoplasmatota* [21-23], also detected in this study but not in the same sampling site.
